# Taylor’s Law, Smith’s Law, and Diversity Power Laws: A Novel Triple Power Law Methodology for Scaling Diversity and Heterogeneity

**DOI:** 10.64898/2026.09.05.749612

**Authors:** Zhanshan (Sam) Ma, Lianwei Li, Aaron M. Ellison

**Affiliations:** Computational Biology and Medical Ecology Lab, Kunming Institute of Zoology, Chinese Academy of Sciences, Kunming, 650221, China; Harvard University, Harvard Forest, 324 North Main Street, Petersham, Massachusetts 01366, USA

**Author notes:** For all correspondence.

**Keywords:** Taylor’s Power Law (TPL), Smith’s Power Law (SPL), Diversity Power law (DPL), Diversity Scaling, Heterogeneity Scaling, Diversity-Heterogeneity Nexus, Diversity-Mean Relationship (DMR), Diversity-Variance Relationship (DVR), Diversity-Heterogeneity Relationship (DHR)

## Abstract

Taylor’s power law (TPL) and Smith’s power law (SPL) are two foundational scaling laws describing how variance scales with mean density and plot area, respectively. While TPL captures ecological heterogeneity (variation among interacting organisms), SPL captures environmental heterogeneity (variation in the abiotic template). Diversity scaling, traditionally approached through species–area relationships, has been extended to diversity–area relationships (DAR) using Hill numbers. Here we integrate TPL, SPL, and DPL, including three newly proposed models [*diversity–mean*, *diversity–variance*, and *diversity–heterogeneity relationship*s (DHR)], into a unified triple power law methodology for scaling diversity and heterogeneity in microbial ecosystems. Using human gut and vaginal microbiome datasets, we systematically vary two orthogonal factors: scale (unit vs. multi-unit) and accrual (without vs. with sample accrual). Our results show that TPL and SPL are complementary: classic TPL captures cross-sectional heterogeneity at the community scale, while *accrual TPL*, a new extension based on sample accrual, captures heterogeneity accumulation at the metacommunity scale and appears less scale-dependent. SPL provides a tool for relating environmental heterogeneity to diversity, supporting the reciprocity principle with ecological heterogeneity. Among the four diversity power laws, DHR, using the variance-to-mean ratio as a direct heterogeneity metric, is most aligned with the diversity–heterogeneity nexus. The triple power law methodology reveals that heterogeneity scaling predicts diversity scaling in the majority of models, with the strongest predictive relationships observed for accrual TPL at higher diversity orders. Nevertheless, the commonly assumed scale-invariance proved elusive, occurring in fewer than 20% of the power law models tested. This framework may extend beyond microbiome ecosystem to any complex system where heterogeneity and diversity arise from interacting components, from ecosystems to economies to artificial intelligence.

## 1. Introduction

Power laws fascinate theoretical ecologists by offering visions of scale invariance and regime-change criticality, but frequently disappoint or confuse practitioners. They often play a key role in quantifying heterogeneity across diverse systems, from market dynamics to ecological dynamics (Heckman, 2001; Cohen et al., 2017; Ji et al., 2019; Shavit & Ellison 2021; Ma & Ellison, 2026; Ma, Liu & Ellison 2026). Taylor’s power law (TPL; Taylor, 1961, Taylor et al. 1977, 1983, 1988) originated in population ecology and later filtered into many fields of science, technology, and even the humanities (Ma & Taylor 2025). It links population mean density (*m*) to its variance (*V*) via (*V* = *am^b^*). Ignoring biological entities, TPL generalizes to a model relating the mean and variance (first and second statistical moments) of any countable random variable. In ecology, it measures the scaling of ecological heterogeneity or temporal stability — a notion extendable to the scaling of heterogeneity and stability in general complex systems (Ma 1991, 2015, 2025). Smith’s power law (SPL; Smith, 1938) relates plot size (S) to crop yield variance via (*V* = *aS*^’*b*^), serving as a measure of environmental heterogeneity. Despite its utility in crop yield study, SPL remains largely unknown outside agricultural science, which is in striking contrast to the ubiquity of TPL (Ma *et al*. 2026).

Both classic SPL (Smith, 1938) and our usage of SPL in this article address the same fundamental question: how does environmental heterogeneity change with scale? They differ in how they operationalize “scale” and “heterogeneity,” yet converge on the same power law form (*V* = *aS*^’*b*^). In classic SPL, scale is represented by plot area (spatial extent), and heterogeneity is measured as the variance of crop yield across replicate plots of the same size. As plot size increases, small-scale environmental variation is averaged out, typically producing a negative scaling exponent. In our usage, scale is represented by the number of accrued samples (sampling effort drawn from the metacommunity), and heterogeneity is measured as the variance of species abundance across species within the accrued composite sample. As more samples are combined, previously unencountered environmental variability is incorporated, producing a positive scaling exponent. Note that classic SPL typically yields a negative exponent (variance decreases with plot size), whereas our formulation produces a positive exponent (variance increases with sampling effort), reflecting the different mechanisms of heterogeneity capture in the two approaches. Despite these operational differences, both formulations capture the scaling of environmental heterogeneity with scale—one through spatial replication across plots, the other through cumulative sampling within a metacommunity. Ma *et al*. (2026) proposed integrating TPL and SPL into a unified framework for scaling ecological and environmental heterogeneity in a reciprocal manner. This unification broadens SPL beyond its agricultural origins, establishing it as a general framework for measuring environmental heterogeneity scaling.

Both classic TPL (Taylor, 1961) and an accrual version of TPL developed in this article both take the form of *V* = *am^b^* and address the same question—how does ecological heterogeneity scale with mean abundance?—but they capture different aspects of this scaling through different sampling designs. Classic TPL operates on independent samples: each sample comes from a single individual in the case of human microbiome study, producing a cloud of (*M*, *V*) pairs that reflect among-individual heterogeneity in species abundance distributions. The scaling exponent *b* thus measures how heterogeneity varies across individuals within the metacommunity of microbial communities. Accrual TPL, by contrast, operates on cumulatively combined samples: as samples are accrued stepwise, both mean abundance and variance increase, tracing a trajectory that reflects how heterogeneity accumulates with sampling effort. The scaling exponent *b* here measures the rate at which heterogeneity builds as more of the metacommunity is sampled. While classic TPL captures a static snapshot of heterogeneity across individuals, accrual TPL captures the dynamic process of heterogeneity accumulation with scale. Both measure ecological heterogeneity, but through different lenses—one cross-sectional, the other cumulative. This distinction is critical: as our results show, the two approaches yield different exponents and exhibit different relationships with other power law models, underscoring that they quantify distinct facets of heterogeneity scaling.

While TPL and SPL address heterogeneity scaling, diversity scaling has traditionally been approached through the species–area relationship (SAR; Watson, 1835). However, species richness is not an ideal metric for diversity, and Hill numbers were introduced to extend SAR into the general diversity–area relationship (DAR; Ma, 2018). DAR takes the form *^q^D* = *cA^z^*, where *^q^D* is the Hill number of order *q* (species richness when *q=0*, Shannon diversity when *q=1*, and Simpson diversity when *q*=2), *A* is area, *c* is a constant, and z is the scaling exponent. In this study, we further extend the traditional DAR model by proposing three additional relationships: diversity–mean (DMR), diversity–variance (DVR), and diversity–heterogeneity (DHR). Collectively, we refer to these four diversity-related power law models as DPL (diversity power laws).

While TPL, SPL, and DPL form the modeling framework for scaling diversity and heterogeneity, scale is a common thread that not only links the three types (families) of power laws but also provides a common testbed for measuring heterogeneity and diversity and comparing their scaling relationships. A key feature of our approach is the explicit distinction between two scales of analysis: the unit scale (US) and the multi-unit scale (MUS). At the US, each sample represents a single individual, and power laws are fitted either across independent individuals (classic TPL) or along an accrual sequence of individual samples (accrual TPL, SPL, DPL). At the MUS, multiple individual samples are randomly combined to form composite samples of size S, where S ranges from 1 to 128.

This scale distinction serves different purposes across the three power law families. For TPL, comparing US and MUS reveals whether ecological heterogeneity scaling is scale-invariant — that is, whether the exponent b remains stable as the sampling unit expands from individuals to increasingly larger aggregates. For SPL, the MUS analysis tests whether environmental heterogeneity scaling depends on the grain size at which the environment is sampled. For DPL, the US-to-MUS transition tests whether diversity scaling with area, mean abundance, variance, and heterogeneity changes when the sampling unit is redefined from individual samples to composite groups — in other words, does diversity scale with these four predictors in the same way regardless of grain size? Across all three model families, the US captures fine-grained, individual-level heterogeneity and diversity patterns, while the MUS reveals emergent scaling properties that arise only when individuals are aggregated into larger units. This dual-scale design thus allows us to distinguish scale-dependent from scale-invariant properties of the diversity–heterogeneity nexus.

As argued by Ma & Ellison (2026), ecological and environmental heterogeneity are reciprocal — a principle that holds significance not only for ecology but also for other complex systems. TPL captures ecological heterogeneity (variation among interacting organisms), while SPL captures environmental heterogeneity (variation in the abiotic template). Together, they form a dual framework for scaling heterogeneity across agents and their environment (Ma et al. 2026). However, diversity (the other fundamental dimension of community structure) has traditionally been studied separately through species–area relationships (SAR) (Watson 1835) and its extensions, DAR (Ma 2018). Integrating heterogeneity scaling (TPL and SPL) (Ma et al. 2026) with diversity power law (DPL) scaling within a unified framework is a logical next step, yet this integration has not been systematically attempted, particularly in microbial ecosystems where both diversity and heterogeneity are exceptionally high.

The objective of this article is to integrate and extend, where necessary, the three power law families (TPL, SPL, and DPL) to investigate the relationship between diversity scaling and heterogeneity scaling, the latter encompassing both ecological and environmental heterogeneity. Specifically, we ask: (i) How does ecological heterogeneity (TPL) scale with mean abundance, and is this scaling scale-invariant or scale-dependent? (ii) How does environmental heterogeneity (SPL) scale with sampling effort, and does it differ between habitats? (iii) How do the four diversity power laws (DAR, DMR, DVR, DHR) scale with their respective predictors, and are they redundant or complementary? (iv) Does heterogeneity scaling predict diversity scaling, and if so, is the relationship positive or negative? (v) Do these scaling relationships differ between the gut and vaginal microbiomes — two contrasting habitats representing high-diversity, even communities versus low-diversity, dominance-driven communities?

To address these questions, we systematically vary two orthogonal factors: scale (unit scale, US, vs. multi-unit scale, MUS) and accrual (without vs. with sample accumulation). This multi-scale, dual-accrual design allows us to distinguish scale-dependent from scale-invariant properties of the diversity–heterogeneity nexus, and to compare cross-sectional (classic) versus cumulative (accrual) modes of heterogeneity measurement. Table 1 outlines the general design; Section 2 details the proposed triple power law methodology, and the subsequent sections present the results and discussion of applying this methodology to two large-scale human microbiome datasets: the American Gut Project (AGP) and the vaginal microbiome. We finally suggest potential links with other key research topics in power law research. By integrating TPL, SPL, and DPL into a single framework, we aim to provide a comprehensive, multi-scale characterization of the diversity–heterogeneity nexus in microbial ecosystems, with implications extending beyond microbiome ecosystems to complex systems in general.

**Table 1.** Triple Power Law Methodology for Scaling Diversity and Heterogeneity.

| Scale (S) | TPL Modeling without Accrual of Sample Units (Fixed Sample Units) | Power Law (PL) Models with Accrual of Sample Units (Variable Sample Units) |  |  |  |  |  |
| --- | --- | --- | --- | --- | --- | --- | --- |
|  |  | TPL (Taylor's power law) | SPL (Smith's power law) | DPL (Diversity Power Law): DAR (Diversity-Area Relation); DMR (Diversity-Mean species abundance Relation), DVR (Diversity-Variance Relation) and DHR (Diversity-Heterogeneity Relation) |  |  |  |
| **Unit Scale (US)** | Classic TPL of US without sample accrual: Section 2.1.0; Table 1, Table S1 | TPL of US with sample accrual: Section 2.1.1; Table 1, Table S1 | SPL of US with sample accrual: Section 2.1.1; Table 1, Table S2 | DAR of US with accrual: Section 2.1.2; Table 2, Table S3 | DMR of US: Section 2.1.2; Table S4 (S=1); Table 1B | DVR of US: Section 2.1.2; Table S4 (S=1); Table 1B | DHR of US: D-(V/M) Section 2.1.2; Table S4 |
| **Multi-Unit Scale (MUS)** | TPL of MUS without sample accrual: Section 2.2.0; Table S1 (Left). | TPL of MUS with sample accrual: Section 2.2.1; Table S1 (Right) | SPL of MUS with accrual: Section 2.2.1; Table S2 | DAR of MUS with accrual: Section 2.2.2; Table S3 | DMR of MUS with accrual: Section 2.2.2; Table S4 | DVR of MUS with accrual: Section 2.2.2; Table S4 | DHR of MUS with accrual: Section 2.2.2; Table S4 |

## 2. The Triple Power Law Method

### 2.0. The Design Principles and Test Datasets

We summarize the proposed method that integrates three families of power laws — including some newly developed models — along two orthogonal design considerations (scale size and sample accrual schemes) (Table 1). These are organized into seven design principles, most of which follow from self-evident first principles.

(i) All power law models — whether for diversity, heterogeneity, or their relationships — take the same mathematical form *y* = *f*(*x*) = *αx^β^*. Nevertheless, while the functional form is identical, the variables *x* and *y* differ across models, and the interpretations of the parameters *α* and *β* carry distinct biological or ecological meanings. We follow traditional conventions for the usages of model parameters (*α* and *β*) and variables (*x* & *y*): for example, *m* for mean, *v* for variance, *z* for diversity scaling and *b* for heterogeneity scaling, a/c for intercept, *m*_0_ for diversity/heterogeneity critical threshold, etc.

(ii) Two primary orthogonal factors — Scale (US vs. MUS) and Sample Accrual (With vs. Without) — divide the power law models into four categories (Table 1). The two categories without accrual are straightforward; these represent classic TPL. Their distinction lies in whether US or MUS is used in fitting the TPL variance–mean (V–M) model, designed to investigate the scale-invariance or scale-dependence of TPL. The two categories with sample accrual use three types of power law models: TPL, SPL, and DPL (Diversity Power Law), each elaborated below.

(iii) TPL measures the scaling of ecological heterogeneity; SPL measures the scaling of environmental heterogeneity; DPL is further divided into four sub-categories: traditional DAR (diversity–area relationship), DMR (diversity–mean species abundance relationship), DVR (diversity–variance relationship), and DHR (diversity–heterogeneity relationship). The last three are newly proposed and are designed to model diversity–heterogeneity relationships.

(iv) Whether TPL, SPL, or DPL, two versions are defined: one with US (the former uses the fixed, basic sampling unit, *e.g*., the microbiome sample of an individual) and one with MUS (the latter uses clusters of unit samples as the modeling scale). All power law models share the same mathematical function, but differ in variables and parameter interpretations.

(v) For SPL modeling at the MUS scale, there are two choices for the independent variable: (a) the natural index number i=1,2,3,…,N (e.g., *N*=1000, the number of resamplings used to build the power law); or (b) the cumulative number of sampling units (quadrats) at each accrual step (*e.g.,* when MUS = 4, the number of sampling units becomes 4, 8, 12, …, 4×1000, when *N*=1000). Both schemes yield the same scaling exponent (b), differing only in the intercept (*a*), which is smaller with the second option.

(vi) Note on HCT (Heterogeneity Critical Threshold): HCT is the value of the independent variable at the tipping point where the dependent variable (heterogeneity) undergoes a phase transition between homogeneous and heterogeneous regimes, marking the random point. For TPL, HCT is the value of mean abundance at this transition. For SPL, HCT is the value of cumulative sample size (or accrual index) at this transition. Below HCT, the system is homogeneous; above HCT, it is heterogeneous.

(vii) A practical consideration for demonstrating and testing the triple power law method is whether sampling is performed with or without replacement. Sampling without replacement is more realistic because no sample is reused, but it requires a large number of data points (samples). Conversely, sampling with replacement is less realistic but can be performed with a limited number of samples, as samples may be reused. We adopt both sampling schemes to fully demonstrate our approach. It is important to note that with either scheme, sampling should be random.

In this article, we use human microbiome datasets, specifically the American Gut Project (AGP; Knight et al., 2019) and a large vaginal microbiome dataset originally published by Doyle et al. (2018), to test the proposed method. The same datasets have been previously reanalyzed for power law modeling (Ma & Taylor, 2020), but with a narrow focus on TPL (Taylor’s power law). In these datasets, one sample was taken from each individual (subject) and is treated as a microbial community (sample).

### 2.1 US (unit-scale) power laws

#### 2.1.0 US-Scale TPL (Taylor power law) Modeling without Sample Accrual

Power law modeling is performed without sample accrual, using each sample as an independent unit at the unit scale (US) — i.e., each sample comes from a single individual. This is simply the classic TPL model. Suppose there are *N* total samples in a microbiome dataset (metacommunity).

(i) For each individual sample *i* (*i*=1,2,…,*N*), compute the mean species size (MSS; *M*_i_), i.e., the mean species abundance across all species in that sample, and its corresponding variance (*V*_i_).

(ii) Retain all *N* pairs of (*M_i_, V_i_*) without combining or accruing any samples.

(iii) Fit Taylor’s power law (TPL) using the *N* pairs of individual sample means (*M_i_*) and variances (*V_i_*). This yields the variance–mean (V–M) power law at the unit scale:

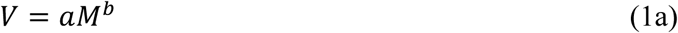

or equivalently,

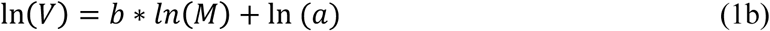

where *M* is the mean species abundance per individual sample, and *V* is the corresponding variance.

Example of classic TPL of US without accrual is displayed in Table 1.

#### 2.1.1 US-Scale TPL and SPL (Smith Power Law) Modeling with Sample Accrual

Power law modeling is performed with accrued samples following the unit scale (US) design — *i.e*., each sample comes from a single individual. Suppose there are *N* total samples in a microbiome dataset (metacommunity).

(i) Compute the mean species size (MSS; *M_1_*), i.e., the mean species abundance across all species in the first community sample, and its corresponding variance (*V*_1_).

(ii) Accrue (combine) the first two samples by summing the abundances of the same species across the two samples. Compute the MSS (*M*_2_) and corresponding variance (*V*_2_) of the combined (accrued) samples.

(iii) Continue accruing the first *S* samples using the same method (*S*=3, 4,…, *N*) and compute the corresponding *M_n_* and *V*_n_ for each accrual step.

(iv) Fit Taylor’s power law (TPL) using the *N* pairs of mean (*M*) and variance (*V*). This yields the variance–mean (V–M) power law:

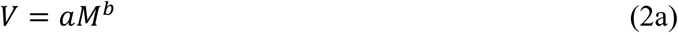

or equivalently,

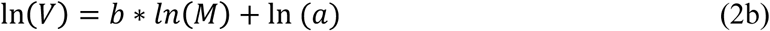

where *M* is the mean of the combined *n* samples, and *V* is the corresponding variance.

(v) Fit Smith’s power law (SPL) using the *N* pairs of accrued sample count (*S*) and variance (*V*). This yields the variance (of MSS)–sample count (V–S) power law:

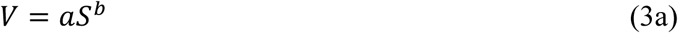

or equivalently,

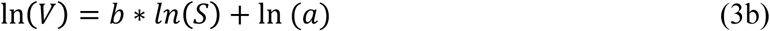

where S is the number of samples accrued at each step, and *V* is the same variance used in fitting Equation (2) in step (iv).

Examples of TPL/SPL fitted at the US scale with sample accrual are illustrated in Table 2 (Table 2A for the sampling scheme with replacement and Table 2B for without replacement).

**Table 2A.**
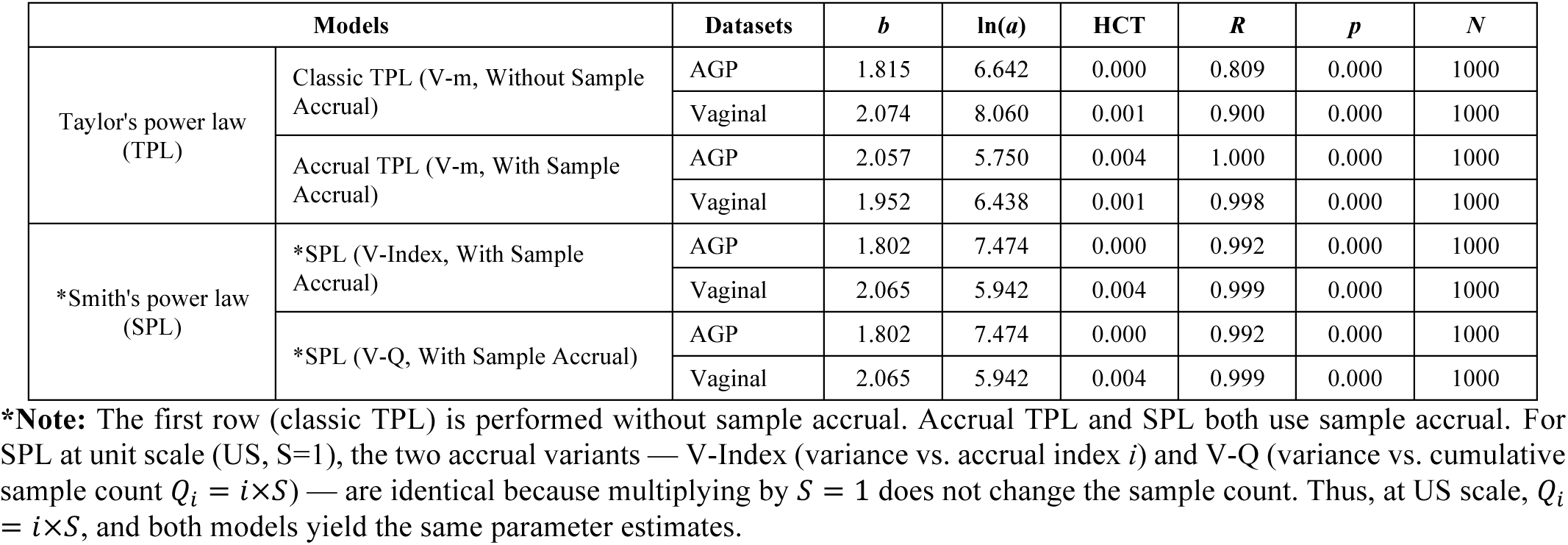
Parameters of Taylor’s power law (TPL) and Smith’s power law (SPL) fitted at the unit scale (US), without and with sample accrual. Results are shown for the American Gut Project (AGP) and Vaginal Microbiome datasets, respectively. Counterpart results fitted at the multi-unit scale (MUS) are reported in Table S1A (for TPL) and Table S2A (for SPL). This Table (2A) is based on the sampling scheme *with replacement* (see Table 2B for results *without replacement*).

| Models | | Datasets | $b$ | $\ln(a)$ | HCT | $R$ | $p$ | $N$ |
| --- | --- | --- | --- | --- | --- | --- | --- | --- |
| Taylor's power law (TPL) | Classic TPL (V-m, Without Sample Accrual) | AGP | 1.815 | 6.642 | 0.000 | 0.809 | 0.000 | 1000 |
|  |  | Vaginal | 2.074 | 8.060 | 0.001 | 0.900 | 0.000 | 1000 |
|  | Accrual TPL (V-m, With Sample Accrual) | AGP | 2.057 | 5.750 | 0.004 | 1.000 | 0.000 | 1000 |
|  |  | Vaginal | 1.952 | 6.438 | 0.001 | 0.998 | 0.000 | 1000 |
| *Smith's power law (SPL) | *SPL (V-Index, With Sample Accrual) | AGP | 1.802 | 7.474 | 0.000 | 0.992 | 0.000 | 1000 |
|  |  | Vaginal | 2.065 | 5.942 | 0.004 | 0.999 | 0.000 | 1000 |
|  | *SPL (V-Q, With Sample Accrual) | AGP | 1.802 | 7.474 | 0.000 | 0.992 | 0.000 | 1000 |
|  |  | Vaginal | 2.065 | 5.942 | 0.004 | 0.999 | 0.000 | 1000 |

**\*Note:** The first row (classic TPL) is performed without sample accrual. Accrual TPL and SPL both use sample accrual. For SPL at unit scale (US, $S=1$ ), the two accrual variants — V-Index (variance vs. accrual index $i$ ) and V-Q (variance vs. cumulative sample count $Q_i = i \times S$ ) — are identical because multiplying by $S = 1$ does not change the sample count. Thus, at US scale, $Q_i = i \times S$ , and both models yield the same parameter estimates.

**Table 2B.**
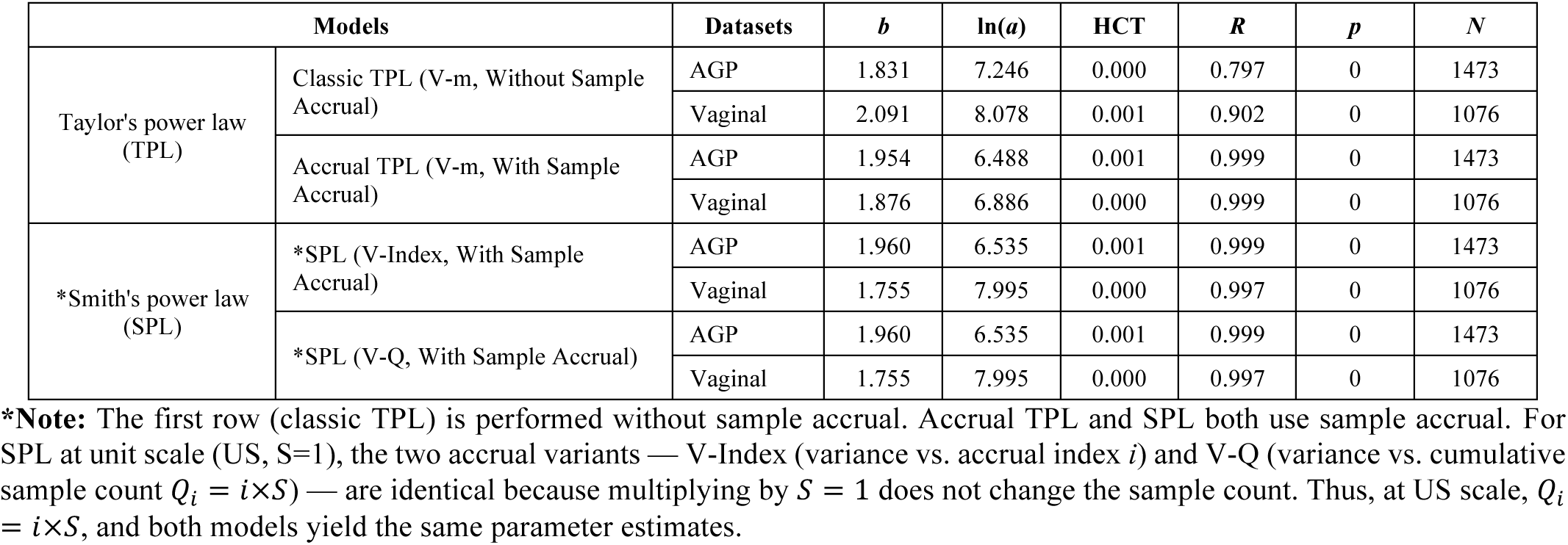
Parameters of Taylor’s power law (TPL) and Smith’s power law (SPL) fitted at the unit scale (US), without and with sample accrual. Results are shown for the American Gut Project (AGP) and Vaginal Microbiome datasets, respectively. Counterpart results fitted at the multi-unit scale (MUS) are reported in Table S1B (for TPL) and Table S2B (for SPL). This Table (2B) is based on the sampling scheme *without replacement* (see Table 2A for results *with replacement*).

#### 2.1.2. US-Scale Diversity Power Law (DPL) Modeling with Sample Accrual

Following the same accrual procedure described in Section 2.1.1 (US-scale with accrual), we compute diversity in Hill numbers at each accrual step *n*=1,2,…,N and fit the following models:

(i) For each accrual step *n*, compute the Hill number diversity (Hill 1973, Chao et al. 2014) *^q^D*(*n*) for orders *q*=0, 1, 2, 3.

(ii) Fit the DAR (Diversity–Area Relationship) model using the accrual step number *n* as the independent variable and *^q^D*(*n*) as the dependent variable: *^q^D* = *cA^z^*. This extends the classic SAR (species–area relationship) to diversity scaling, i.e.,

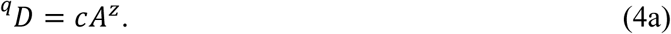

(iii) Fit the DMR (Diversity–Mean species abundance Relationship) model using the mean species abundance *M_n_* (computed in Section 2.1.1) as the independent variable and *^q^D*(*n*) as the dependent variable. The new DMR model we propose is:

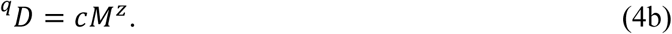

(iv) Fit the DVR (Diversity–Variance Relationship) model using the variance *V_n_* (computed in Section 2.1.1) as the independent variable and *^q^D*(*n*) as the dependent variable, *^q^D* = *cV^z^*, following the same fitting procedure as in (iii), i.e.,

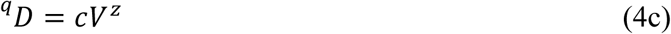

(v) Fit the DHR (Diversity-Heterogeneity Relationship) model using the *H_n_*=*V_n_/M_n_* (computed in Section 2.1.1) as the independent variable and *^q^D*(*n*) as the dependent variable, *^q^D* = *cH^z^*, following the same fitting procedure as in (iii), i.e.,

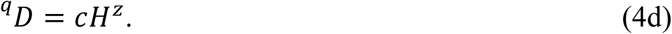

Examples of diversity power law (DPL) scaling fitted at the US scale with sample accrual are illustrated in Table 3 (Table 3A for the sampling scheme with replacement and Table 3B for without replacement).

**Table 3A.** Parameters of four diversity power law models [DAR (diversity-area relationship), DMR (diversity-mean relationship), DVR (diversity-variance relationship), and DHR (diversity-heterogeneity relationship) at unit scale (US, S=1). The counterpart results fitted at multi-unit scale (MUS) are reported in Table S3A (for DAR) and Table S4A (for DMR, DVR, and DHR). All diversity power law modes are built with sample accrual. This Table 3A is based on the sampling scheme *with replacement* (see Table 3B for resu<u>lts *without replacement*).</u>

| Models | Datasets | Orders | $b$ | $\ln(a)$ | $R$ | $p$ | $N$ |
| --- | --- | --- | --- | --- | --- | --- | --- |
| DAR (diversity-area relationship):<br>(D-Index)<br>Classic DAR | AGP (American Gut Microbiome Project) | $q=0$ | 0.254 | 7.426 | 0.996 | 0.000 | 1000 |
| | | $q=1$ | 0.056 | 5.066 | 0.805 | 0.000 | 1000 |
| | | $q=2$ | -0.046 | 4.206 | 0.624 | 0.000 | 1000 |
| | | $q=3$ | -0.083 | 3.796 | 0.806 | 0.000 | 1000 |
| | Vaginal Microbiome | $q=0$ | 0.757 | 3.970 | 0.998 | 0.000 | 1000 |
| | | $q=1$ | 0.135 | 3.540 | 0.775 | 0.000 | 1000 |
| | | $q=2$ | 0.048 | 3.134 | 0.344 | 0.000 | 1000 |
| | | $q=3$ | 0.017 | 2.961 | 0.116 | 0.000 | 1000 |
| DAR (D-Q) | **Note:** At US scale ($S=1$), accrual index $i$ and cumulative sample count $Q_i$ are identical ($Q_i=i$), so DAR/DMR/DVR/DHR parameter estimates are the same for both variants. | | | | | | |
| DMR (diversity-mean relationship):<br>(D-m) | AGP | $q=0$ | 0.285 | 7.213 | 0.987 | 0.000 | 1000 |
| | | $q=1$ | 0.059 | 5.043 | 0.747 | 0.000 | 1000 |
| | | $q=2$ | -0.057 | 4.279 | 0.688 | 0.000 | 1000 |
| | | $q=3$ | -0.098 | 3.899 | 0.846 | 0.000 | 1000 |
| | Vaginal | $q=0$ | 0.716 | 4.146 | 0.999 | 0.000 | 1000 |
| | | $q=1$ | 0.129 | 3.560 | 0.788 | 0.000 | 1000 |
| | | $q=2$ | 0.048 | 3.132 | 0.361 | 0.000 | 1000 |
| | | $q=3$ | 0.018 | 2.952 | 0.132 | 0.000 | 1000 |
| DVR (diversity-variance relationship):<br>(D-V) | AGP | $q=0$ | 0.138 | 6.428 | 0.984 | 0.000 | 1000 |
| | | $q=1$ | 0.028 | 4.889 | 0.730 | 0.000 | 1000 |
| | | $q=2$ | -0.028 | 4.453 | 0.709 | 0.000 | 1000 |
| | | $q=3$ | -0.049 | 4.188 | 0.860 | 0.000 | 1000 |
| | Vaginal | $q=0$ | 0.365 | 1.821 | 0.995 | 0.000 | 1000 |
| | | $q=1$ | 0.063 | 3.196 | 0.748 | 0.000 | 1000 |
| | | $q=2$ | 0.020 | 3.049 | 0.302 | 0.000 | 1000 |
| | | $q=3$ | 0.005 | 2.971 | 0.070 | 0.026 | 1000 |
| DHR (diversity-heterogeneity relationship):<br>(D-H) | AGP | $q=0$ | 0.267 | 5.695 | 0.979 | 0.000 | 1000 |
| | | $q=1$ | 0.053 | 4.752 | 0.713 | 0.000 | 1000 |
| | | $q=2$ | -0.057 | 4.625 | 0.728 | 0.000 | 1000 |
| | | $q=3$ | -0.096 | 4.468 | 0.872 | 0.000 | 1000 |
| | Vaginal | $q=0$ | 0.737 | -0.518 | 0.987 | 0.000 | 1000 |
| | | $q=1$ | 0.120 | 2.872 | 0.705 | 0.000 | 1000 |
| | | $q=2$ | 0.033 | 3.020 | 0.239 | 0.000 | 1000 |
| | | $q=3$ | 0.001 | 3.051 | 0.006 | 0.857 | 1000 |

**Table 3B.** Parameters of four diversity power law models [DAR (diversity-area relationship), DMR (diversity-mean relationship), DVR (diversity-variance relationship), and DHR (diversity-heterogeneity relationship) at unit scale (US, S=1). The counterpart results fitted at multi-unit scale (MUS) are reported in Table S3B (for DAR) and Table S4B (for DMR, DVR, and DHR). All diversity power law modes are built with sample accrual. This Table 3B is based on the sampling scheme *with replacement* (see Table 3A for results *w<u>ithout replacement</u>*).

| Models | Datasets | Orders | $b$ | $\ln(a)$ | $R$ | $p$ | $N$ |
| --- | --- | --- | --- | --- | --- | --- | --- |
| DAR (diversity-area relationship):<br>(D-Index)<br>Classic DAR | AGP (American Gut Microbiome Project) | $q=0$ | 0.298 | 7.169 | 0.987 | 0.000 | 1473 |
| | | $q=1$ | 0.077 | 4.897 | 0.679 | 0.000 | 1473 |
| | | $q=2$ | 0.046 | 3.544 | 0.466 | 0.000 | 1473 |
| | | $q=3$ | 0.037 | 2.946 | 0.433 | 0.000 | 1473 |
| | Vaginal Microbiome | $q=0$ | 0.811 | 3.882 | 0.998 | 0.000 | 1076 |
| | | $q=1$ | 0.204 | 3.109 | 0.907 | 0.000 | 1076 |
| | | $q=2$ | 0.115 | 2.762 | 0.780 | 0.000 | 1076 |
| | | $q=3$ | 0.077 | 2.683 | 0.632 | 0.000 | 1076 |
| DAR (diversity-area relationship): (D-Q) | AGP (American Gut Microbiome Project) | $q=0$ | 0.298 | 7.169 | 0.987 | 0.000 | 1473 |
| | | $q=1$ | 0.077 | 4.897 | 0.679 | 0.000 | 1473 |
| | | $q=2$ | 0.046 | 3.544 | 0.466 | 0.000 | 1473 |
| | | $q=3$ | 0.037 | 2.946 | 0.433 | 0.000 | 1473 |
| | Vaginal Microbiome | $q=0$ | 0.811 | 3.882 | 0.998 | 0.000 | 1076 |
| | | $q=1$ | 0.204 | 3.109 | 0.907 | 0.000 | 1076 |
| | | $q=2$ | 0.115 | 2.762 | 0.780 | 0.000 | 1076 |
| | | $q=3$ | 0.077 | 2.683 | 0.632 | 0.000 | 1076 |
| DMR (diversity-mean relationship):<br>(D-m) | AGP | $q=0$ | 0.297 | 7.159 | 0.989 | 0.000 | 1473 |
| | | $q=1$ | 0.078 | 4.893 | 0.683 | 0.000 | 1473 |
| | | $q=2$ | 0.046 | 3.543 | 0.467 | 0.000 | 1473 |
| | | $q=3$ | 0.037 | 2.946 | 0.432 | 0.000 | 1473 |
| | Vaginal | $q=0$ | 0.863 | 3.392 | 0.996 | 0.000 | 1076 |
| | | $q=1$ | 0.217 | 2.983 | 0.906 | 0.000 | 1076 |
| | | $q=2$ | 0.124 | 2.685 | 0.786 | 0.000 | 1076 |
| | | $q=3$ | 0.084 | 2.623 | 0.647 | 0.000 | 1076 |
| DVR (diversity-variance relationship):<br>(D-V) | AGP | $q=0$ | 0.151 | 6.194 | 0.982 | 0.000 | 1473 |
| | | $q=1$ | 0.038 | 4.672 | 0.651 | 0.000 | 1473 |
| | | $q=2$ | 0.022 | 3.428 | 0.427 | 0.000 | 1473 |
| | | $q=3$ | 0.017 | 2.856 | 0.391 | 0.000 | 1473 |
| | Vaginal | $q=0$ | 0.458 | 0.262 | 0.993 | 0.000 | 1076 |
| | | $q=1$ | 0.113 | 2.241 | 0.883 | 0.000 | 1076 |
| | | $q=2$ | 0.063 | 2.284 | 0.753 | 0.000 | 1076 |
| | | $q=3$ | 0.042 | 2.368 | 0.608 | 0.000 | 1076 |
| DHR (diversity-heterogeneity relationship):<br>(D-H) | AGP | $q=0$ | 0.305 | 5.216 | 0.973 | 0.000 | 1473 |
| | | $q=1$ | 0.073 | 4.470 | 0.615 | 0.000 | 1473 |
| | | $q=2$ | 0.040 | 3.338 | 0.384 | 0.000 | 1473 |
| | | $q=3$ | 0.031 | 2.789 | 0.348 | 0.000 | 1473 |
| | Vaginal | $q=0$ | 0.970 | -3.201 | 0.986 | 0.000 | 1076 |
| | | $q=1$ | 0.233 | 1.463 | 0.855 | 0.000 | 1076 |
| | | $q=2$ | 0.127 | 1.884 | 0.712 | 0.000 | 1076 |
| | | $q=3$ | 0.082 | 2.128 | 0.561 | 0.000 | 1076 |

### 2.2. Multi-Unit Scale (MUS) Power Laws

#### 2.2.0. MUS-Scale TPL Modeling Without Sample Accrual

This extends the classic TPL from unit scale (US) to multi-unit scale (MUS) by randomly combining multiple unit samples (each from a single individual) into larger composite samples.

While classic TPL (Section 2.1.0) is applied at the unit scale — i.e., each sample comes from a single individual — the multi-unit scale (MUS) TPL extends the analysis to composite samples formed by randomly combining multiple unit samples. This allows us to investigate scale-invariance theory whether TPL scaling parameters change as the sampling unit size (number of individuals per composite sample) increases.

Importantly, like classic TPL, this MUS variant is also without sample accrual — each composite sample is treated as an independent unit, and samples are not cumulatively combined across steps. The procedure is analogous to classic TPL but operates on aggregated samples rather than individual ones.

(i) Randomly select *S* unit samples (with replacement) from the dataset of *N* total samples. Combine these *S* samples by summing the abundances of the same species across samples. Compute the mean species abundance (M) and the corresponding variance (V) across all species in the combined sample.

(ii) Repeat step (i) 1000 times for a fixed S to obtain 1000 pairs of mean (*M*) and variance (*V*). Fit Taylor’s power law (*V* = *aM^b^*) to these 1000 pairs.

(iii) Repeat steps (i) and (ii) for a series of MUS a full spectrum *S*=1, 2, 3,…, 128.

The results from fitting the TPL at MUS scale is presented in Table S1 (the left section).

#### 2.2.1. MUS-Scale TPL and SPL Modeling with Sample Accrual

This section extends the accrual procedure from unit scale (US) to multi-unit scale (MUS). Unlike MUS without accrual (Section 2.2.0), where each composite sample is independent, this method cumulatively accrues samples, analogous to the US accrual procedure described in Section 2.1.1.

(i) Randomly select *S* unit samples from the dataset. Combine these *S* samples by summing the abundances of the same species across samples to create a new composite sample *C_1_*. Compute the variance (V) across all species in *C_1_*. Record the accrual index as *i*=1 and the cumulative sample count (scale size) as Q=S.

(ii) Randomly select another *S* unit samples (with replacement) from the dataset. Combine these *S* samples with *C_1_* to create a new composite sample *C_2_*. Compute the variance (V) across all species in C_2_. Record the accrual index as i=2 and the cumulative sample count as Q=2S.

(iii) Repeat step (ii) iteratively: at each iteration *i*, combine a newly selected set of *S* unit samples with the composite sample from the previous iteration to form a new composite sample *C*_i._ Compute the variance (*V*) for each *C_i_*. Increment the accrual index by 1 and increase the cumulative sample count by S each time. Continue until the accrual index reaches *i*=1000.

(iv-A) Fit TPL (Taylor power law model) with accrual: variance *V vs*. mean species abundance *m*. The results for S=1-128 are exhibited in the right section of table S1.

(iv-B) Fit two SPL power law models: (*a*) variance *V* vs. accrual index *i*, and (*b*) variance *V* vs. cumulative sample count Q. The former captures scaling with iteration number; the latter captures scaling with total sample size. The results for S=1-128 are exhibited in the right section of Table S2.

#### 2.2.2. MUS-Scale Diversity Power Law (DPL) Modeling

Following the same accrual procedure described in Section 2.2.1 (MUS-scale with accrual), we compute diversity in Hill numbers at each accrual step and fit the following models. This extends the US-scale diversity models (Section 2.1.2) to multi-unit scale, where each accrual step combines multiple unit samples rather than single individuals.

(i) For each accrual step *i*=1,2,…,1000 and cumulative sample count *Q_i_*=*i*×*S* (where *S* is the fixed MUS size), compute the Hill number diversity *^q^D*(*i*) for orders *q* = 0, 1, 2, 3.

(ii) Fit the DAR (Diversity–Area Relationship) model at MUS scale using the accrual index *i* as the independent variable and *^q^D*(*i*) as the dependent variable. This is the MUS-scale counterpart of the US-scale DAR. Results for the DAR are presented in Table S3.

(iii) Fit the DMR (Diversity–Mean species abundance Relationship) model at MUS scale using the mean species abundance *M_i_* (computed from the accrued composite sample at step *i*) as the independent variable and *^q^D*(*i*) as the dependent variable. This is the MUS-scale version of DMR.

(iv) Fit the DVR (Diversity–Variance Relationship) model at MUS scale using the variance V_i_ (computed from the accrued composite sample at step *i*) as the independent variable and *^q^D*(*i*) as the dependent variable, following the same fitting procedure as in (iii). This is the MUS-scale version of DVR.

(v) Fit the DHR (Diversity-Heterogeneity Relationship) model at MUS scale using the *H_i_*=*V_i_/M_i_* (computed in Section 2.2.1) as the independent variable and *^q^D*(*i*) as the dependent variable, *^q^D* = *cH^z^*, following the same fitting procedure as in (iii).

Results for DMR, DVR and DHR models at MUS scale are presented in Table S4, while those for DAR are in Table S3 as mentioned previously. Table S5 presented the Spearman’s correlations between diversity, mean species abundance, and variance, as well as heterogeneity (V/m), which further verify the validities of newly proposed DPL (diversity power law) models.

### 2.3 Statistical Correlation Analyses between the Three Power Law families

#### 2.3.0 Correlation Analysis between Classic TPL, Accrual TPL, and SPL Across MUS Scales

To systematically compare parameter estimates across different power law models, we first calculated Spearman’s rank correlations between counterpart parameters. Note that all SPL models are inherently accrual-based (variance scales with cumulative sample count or accrual index), whereas TPL can be implemented either with accrual (variance vs. mean of accrued samples) or without accrual (classic TPL: variance vs. mean of independent samples). Comparisons were designed to assess: (i) whether classic TPL (non-accrual) correlates with accrual-based SPL and TPL; (ii) whether different accrual variants of SPL (Q-V vs. Index-V) are equivalent; and (iii) whether SPL variants correlate with accrual TPL. All six pairwise comparisons (between classic TPL, accrual TPL, SPL Q-V, and SPL Index-V) were performed separately for gut and vaginal microbiomes across MUS scales (*S* = 1 *to* 128), and the results are summarized in Table S6.

#### 2.3.1 Correlation Analysis between Classic DAR and New Diversity Power Laws (DPLs)

Next, we performed pairwise Spearman’s rank correlations between the four DPL models, including traditional DAR and the three new DPLs, *i.e*., DMR (diversity–mean relationship), DVR (diversity–variance relationship), and DHR (diversity–heterogeneity relationship). To simplify the comparison, we only compared the scaling parameter (*z*). A total of six pairwise comparisons were performed for each microbiome habitat (AGP and vaginal) and for each diversity order *q*. The results are summarized in Table S6.

#### 2.3.2 Correlation Analysis between Classic TPL, Accrual TPL, SPL, and Diversity Power Laws

We also compared classic TPL, accrual TPL, and SPL with the four DPL models by correlating their scaling parameters (*b* for TPL/SPL *vs*. *z* for DPLs) for each diversity order *q.* These correlations were performed separately for AGP and vaginal datasets using Spearman’s rank correlation coefficients (Table S6).

### 2.4 Habitat Comparison of Power Law Parameters

To test whether power law parameters differ between gut (AGP) and vaginal microbiomes, we performed Wilcoxon rank-sum tests for each parameter [b, ln(a), HCT] across four heterogeneity model types: classic TPL without accrual (m-V Non-Accrual), accrual TPL (m-V Accrual), and two SPL accrual variants (Q-V Accrual, Index-V Accrual). All comparisons were based on MUS scale results across *S* = 1 *to* 128 (Table S7).

We further tested differences between AGP and vaginal microbiomes for the corresponding parameters of the four DPL models — DAR, DMR, DVR, and DHR — at each diversity order q. The results are also summarized in Table S7.

### 2.5. Bridging the Three Families of Power Laws: TPL, SPL, and DPLs

The preceding sections (2.3 and 2.4) established the foundations for this subsection by examining pairwise correlations and habitat differences among the three power law families. Here, we close the gap by establishing direct statistical models between their scaling parameters. As it turned out, most relationships between the scaling parameters of TPL, SPL, and DPL models can be adequately described by simple linear regression models of the form *y* = *a* + *bx*.

To further quantify these relationships, we performed linear regression analyses between the scaling parameters of TPL, SPL, and DPL models. Table 3 presents the regression parameters for *b*-values between classic TPL, accrual TPL, and SPL (both V-i and V-Q variants). Table 2 presents the regression parameters for z values between DAR (D-Q), DMR, DVR, and DHR across diversity orders. Table 3 presents the regression parameters between *b*-values of TPL/SPL models and *z*-values of DPL models.

**Table 4.** Parameters of linear regression models relating the *b*-values of classic TPL, accrual TPL, SPL (V-i), and SPL (V-Q) pairwise, with microbiome sites (AGP or vaginal) and sampling schemes (with/without replacement) all modeled separately (four regimes in total).

| Relation Model<br>$b_1 = \alpha + \beta b_2$ | AGP | | | | Vaginal Microbiome | | | | $N$ |
| --- | --- | --- | --- | --- | --- | --- | --- | --- | --- |
| | $\beta$ | $\alpha$ | $R$ | $p$ | $\beta$ | $\alpha$ | $R$ | $p$ | |
| With Replacement Sampling |  |  |  |  |  |  |  |  |  |
| Classic TPL vs. SPL(i) | -0.001 | 1.999 | 0.002 | 0.980 | 0.052 | 1.875 | 0.141 | 0.112 | 128 |
| Classic TPL vs. SPL(Q) | -0.001 | 1.999 | 0.002 | 0.980 | 0.052 | 1.875 | 0.141 | 0.112 | 128 |
| Classic TPL vs. Accrual TPL | -0.034 | 2.063 | 0.252 | 0.004 | 0.076 | 1.820 | 0.260 | 0.003 | 128 |
| SPL(i) vs. SPL(Q) | 1.000 | 0.000 | 1.000 | 0.000 | 1.000 | 0.000 | 1.000 | 0.000 | 128 |
| SPL(i) vs. Accrual TPL | -0.104 | 2.206 | 0.247 | 0.005 | 0.628 | 0.741 | 0.790 | 0.000 | 128 |
| SPL(Q) vs. Accrual TPL | -0.104 | 2.206 | 0.247 | 0.005 | 0.628 | 0.741 | 0.790 | 0.000 | 128 |
| Without Replacement Sampling |  |  |  |  |  |  |  |  |  |
| Classic TPL vs. SPL(i) | 0.094 | 1.793 | 0.686 | 0.003 | -0.048 | 1.856 | 0.651 | 0.006 | 16 |
| Classic TPL vs. SPL(Q) | 0.094 | 1.793 | 0.686 | 0.003 | -0.048 | 1.856 | 0.651 | 0.006 | 16 |
| Classic TPL vs. Accrual TPL | 0.108 | 1.758 | 0.648 | 0.007 | -0.041 | 1.978 | 0.285 | 0.285 | 16 |
| SPL(i) vs. SPL(Q) | 1.000 | 0.000 | 1.000 | 0.000 | 1.000 | 0.000 | 1.000 | 0.000 | 16 |
| SPL(i) vs. Accrual TPL | 1.168 | -0.339 | 0.962 | 0.000 | -0.013 | 1.916 | 0.007 | 0.981 | 16 |
| SPL(Q) vs. Accrual TPL | 1.168 | -0.339 | 0.962 | 0.000 | -0.013 | 1.916 | 0.007 | 0.981 | 16 |

**Table 5.** Parameters of linear regression models relating the z-values of DAR (D-Q), DMR, DVR, and DHR pairwise, with microbiome sites (AGP or vaginal) and sampling schemes (with/without replacement) all modeled separately (four regimes in total).

| Diversity Order | Relation Model<br>$z_1 = \alpha + \beta z_2$ | AGP | | | | Vaginal Microbiome | | | | N |
| --- | --- | --- | --- | --- | --- | --- | --- | --- | --- | --- |
| | | $\beta$ | $\alpha$ | R | p | $\beta$ | $\alpha$ | R | p | |
| **With Replacement** |  |  |  |  |  |  |  |  |  |  |
| $q=0$ | DAR(Q) vs. DMR | 1.016 | -0.001 | 0.998 | 0.000 | 0.989 | 0.001 | 1.000 | 0.000 | 128 |
|  | DAR(Q) vs. DVR | 0.505 | 0.000 | 0.999 | 0.000 | 0.508 | -0.001 | 0.999 | 0.000 | 128 |
|  | DAR(Q) vs. DHR | 1.003 | 0.000 | 0.999 | 0.000 | 1.041 | -0.004 | 0.997 | 0.000 | 128 |
|  | DMR vs. DVR | 0.497 | 0.000 | 1.000 | 0.000 | 0.514 | -0.001 | 1.000 | 0.000 | 128 |
|  | DMR vs. DHR | 0.986 | 0.001 | 0.999 | 0.000 | 1.053 | -0.006 | 0.998 | 0.000 | 128 |
|  | DVR vs. DHR | 1.986 | 0.000 | 1.000 | 0.000 | 2.052 | -0.003 | 1.000 | 0.000 | 128 |
| $q=1$ | DAR(Q) vs. DMR | 1.007 | 0.000 | 1.000 | 0.000 | 0.999 | 0.000 | 1.000 | 0.000 | 128 |
|  | DAR(Q) vs. DVR | 0.496 | 0.000 | 1.000 | 0.000 | 0.507 | 0.000 | 0.998 | 0.000 | 128 |
|  | DAR(Q) vs. DHR | 0.978 | 0.000 | 0.999 | 0.000 | 1.024 | -0.001 | 0.996 | 0.000 | 128 |
|  | DMR vs. DVR | 0.493 | 0.000 | 1.000 | 0.000 | 0.508 | 0.000 | 0.999 | 0.000 | 128 |
|  | DMR vs. DHR | 0.970 | 0.000 | 0.999 | 0.000 | 1.027 | -0.001 | 0.998 | 0.000 | 128 |
|  | DVR vs. DHR | 1.971 | 0.000 | 1.000 | 0.000 | 2.024 | 0.000 | 0.999 | 0.000 | 128 |
| $q=2$ | DAR(Q) vs. DMR | 1.056 | 0.000 | 0.995 | 0.000 | 1.011 | 0.000 | 1.000 | 0.000 | 128 |
|  | DAR(Q) vs. DVR | 0.526 | 0.000 | 0.995 | 0.000 | 0.513 | 0.000 | 0.998 | 0.000 | 128 |
|  | DAR(Q) vs. DHR | 1.048 | 0.000 | 0.995 | 0.000 | 1.038 | -0.001 | 0.995 | 0.000 | 128 |
|  | DMR vs. DVR | 0.499 | 0.000 | 1.000 | 0.000 | 0.508 | 0.000 | 0.999 | 0.000 | 128 |
|  | DMR vs. DHR | 0.993 | 0.000 | 0.999 | 0.000 | 1.028 | -0.001 | 0.996 | 0.000 | 128 |
|  | DVR vs. DHR | 1.992 | 0.000 | 1.000 | 0.000 | 2.027 | 0.000 | 0.999 | 0.000 | 128 |
| $q=3$ | DAR(Q) vs. DMR | 1.093 | 0.000 | 0.996 | 0.000 | 1.010 | 0.000 | 1.000 | 0.000 | 128 |
|  | DAR(Q) vs. DVR | 0.543 | 0.000 | 0.997 | 0.000 | 0.515 | 0.000 | 0.999 | 0.000 | 128 |
|  | DAR(Q) vs. DHR | 1.076 | 0.000 | 0.997 | 0.000 | 1.048 | -0.001 | 0.995 | 0.000 | 128 |
|  | DMR vs. DVR | 0.496 | 0.000 | 1.000 | 0.000 | 0.510 | 0.000 | 0.999 | 0.000 | 128 |
|  | DMR vs. DHR | 0.984 | 0.000 | 1.000 | 0.000 | 1.037 | -0.001 | 0.996 | 0.000 | 128 |
|  | DVR vs. DHR | 1.983 | 0.000 | 1.000 | 0.000 | 2.038 | 0.000 | 0.999 | 0.000 | 128 |
| **Without Replacement** |  |  |  |  |  |  |  |  |  |  |
| $q=0$ | DAR(Q) vs. DMR | 0.964 | 0.010 | 0.999 | 0.000 | 0.485 | 0.473 | 0.347 | 0.188 | 16 |
|  | DAR(Q) vs. DVR | 0.572 | -0.020 | 0.998 | 0.000 | 0.699 | -0.109 | 0.901 | 0.000 | 16 |
|  | DAR(Q) vs. DHR | 1.264 | -0.071 | 0.996 | 0.000 | 2.399 | -0.980 | 0.888 | 0.000 | 16 |
|  | DMR vs. DVR | 0.594 | -0.025 | 0.999 | 0.000 | 0.319 | 0.179 | 0.576 | 0.020 | 16 |
|  | DMR vs. DHR | 1.311 | -0.084 | 0.997 | 0.000 | 0.187 | 0.793 | 0.097 | 0.722 | 16 |
|  | DVR vs. DHR | 2.211 | -0.028 | 0.999 | 0.000 | 3.021 | -0.419 | 0.868 | 0.000 | 16 |
| $q=1$ | DAR(Q) vs. DMR | 1.017 | -0.001 | 1.000 | 0.000 | 1.049 | 0.003 | 0.997 | 0.000 | 16 |
|  | DAR(Q) vs. DVR | 0.478 | 0.001 | 1.000 | 0.000 | 0.596 | -0.009 | 0.998 | 0.000 | 16 |
|  | DAR(Q) vs. DHR | 0.878 | 0.005 | 0.999 | 0.000 | 1.329 | -0.037 | 0.995 | 0.000 | 16 |
|  | DMR vs. DVR | 0.470 | 0.001 | 1.000 | 0.000 | 0.566 | -0.010 | 0.998 | 0.000 | 16 |
|  | DMR vs. DHR | 0.863 | 0.006 | 0.998 | 0.000 | 1.258 | -0.039 | 0.991 | 0.000 | 16 |
|  | DVR vs. DHR | 1.838 | 0.003 | 0.999 | 0.000 | 2.231 | -0.018 | 0.998 | 0.000 | 16 |
| $q=2$ | DAR(Q) vs. DMR | 1.022 | -0.001 | 1.000 | 0.000 | 1.092 | -0.002 | 0.999 | 0.000 | 16 |
|  | DAR(Q) vs. DVR | 0.474 | 0.000 | 1.000 | 0.000 | 0.590 | -0.005 | 0.999 | 0.000 | 16 |
|  | DAR(Q) vs. DHR | 0.859 | 0.000 | 0.998 | 0.000 | 1.257 | -0.016 | 0.996 | 0.000 | 16 |
|  | DMR vs. DVR | 0.463 | 0.000 | 1.000 | 0.000 | 0.540 | -0.004 | 0.999 | 0.000 | 16 |
|  | DMR vs. DHR | 0.840 | 0.000 | 0.998 | 0.000 | 1.150 | -0.014 | 0.995 | 0.000 | 16 |
|  | DVR vs. DHR | 1.815 | 0.000 | 0.999 | 0.000 | 2.133 | -0.007 | 0.999 | 0.000 | 16 |
| $q=3$ | DAR(Q) vs. DMR | 1.022 | -0.001 | 1.000 | 0.000 | 1.114 | -0.002 | 1.000 | 0.000 | 16 |
|  | DAR(Q) vs. DVR | 0.474 | 0.000 | 1.000 | 0.000 | 0.585 | -0.003 | 0.999 | 0.000 | 16 |
|  | DAR(Q) vs. DHR | 0.862 | -0.001 | 0.998 | 0.000 | 1.211 | -0.009 | 0.996 | 0.000 | 16 |
|  | DMR vs. DVR | 0.464 | 0.000 | 1.000 | 0.000 | 0.525 | -0.002 | 0.999 | 0.000 | 16 |
|  | DMR vs. DHR | 0.843 | 0.000 | 0.997 | 0.000 | 1.086 | -0.007 | 0.996 | 0.000 | 16 |
|  | DVR vs. DHR | 1.818 | 0.000 | 0.999 | 0.000 | 2.072 | -0.004 | 0.999 | 0.000 | 16 |

All regressions were performed separately for AGP and vaginal microbiomes, using parameters estimated from MUS scales with and without replacement sampling, respectively. For each regression, we report the slope (*b*), intercept [ln(*a*)], linear correlation coefficient (*R*), and associated *p*-value.

## 3. Application to Human Microbiome Ecosystems

All analyses are performed under two different sampling schemes: without and with replacement. Given that both sets of results exhibit the same or similar patterns, our interpretation and discussion are based on the sampling scheme with replacement. Tables with replacement are labeled A (e.g., Table 2A, Table S1A), and those without replacement are labeled B (e.g., Table 2B, Table S1B). Figures 1–5 present results from modeling the AGP dataset under the sampling scheme with replacement; corresponding results for the vaginal microbiome dataset are provided in the Online Supplementary Information (OSI). Specifically, Figure 1 (A–D) shows the fittings of eight power law models: classic TPL, new accrual TPL, two SPL variants (V-Index and V-Q), and four DPL (DAR, DMR, DVR, and DHR) models. Figure 2 (A–B) displays the relationships between TPL/SPL parameters and scale size (*S* = 1 *to* 128). Figure 3 (A–D) shows the relationships between DPL parameters and scale size (*S* = 1 *to* 128) for each diversity order (*q* = 0 *to* 3). Figure 4 (A–I) presents Spearman’s correlations of scaling exponents between different power law models across scale sizes. Figure 5 illustrates an example of scale invariance testing based on the AGP dataset.

**figure 1A.**
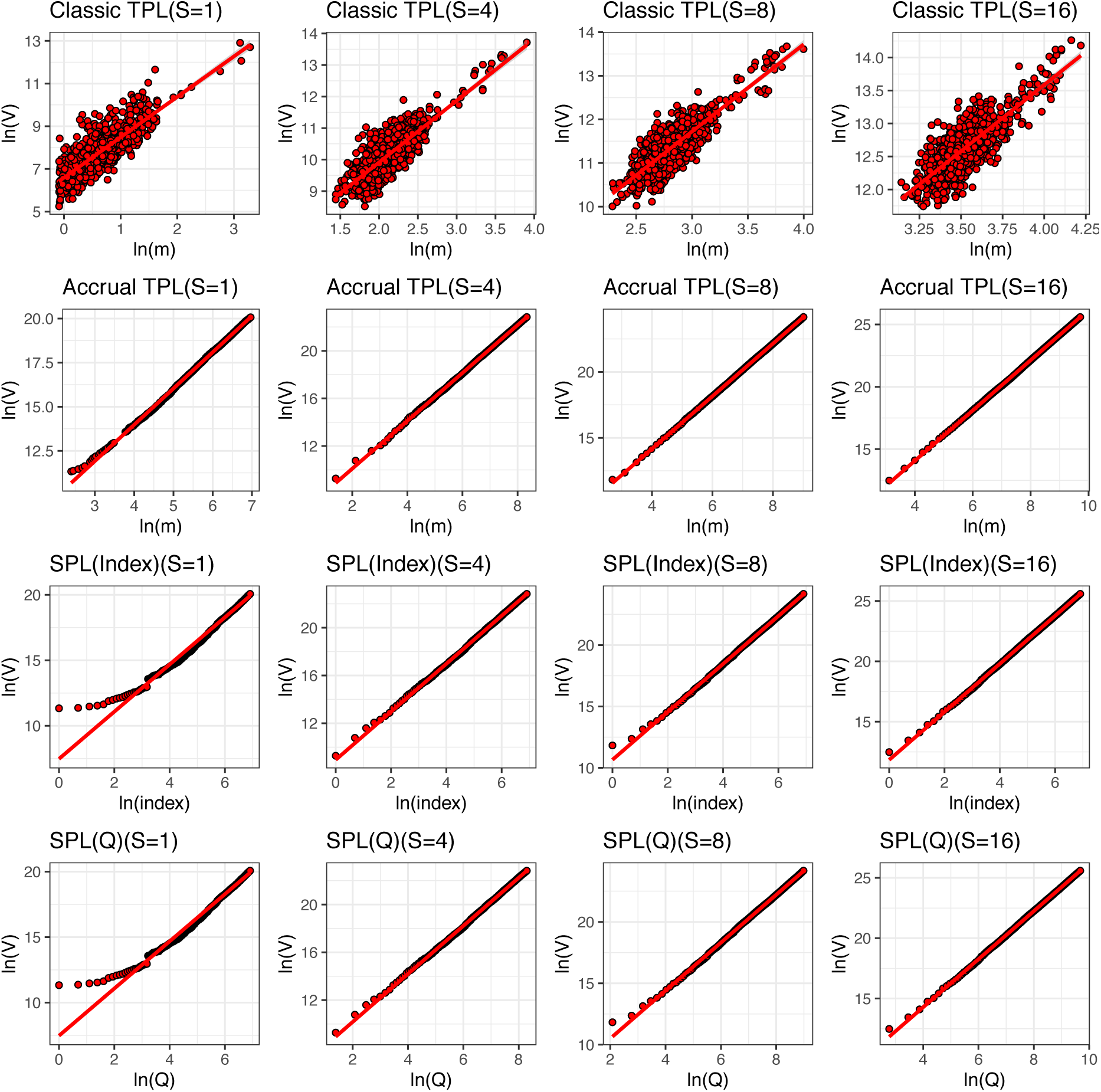
Classic TPL (Taylor’s power law, without sample accrual), new TPL *with sample accrual*, and classic SPL (Smith’s power law) fitted to the AGP (American Gut Project) datasets at MUS (multi-unit scale) scales *S* = 1, 4, 8, 16. The *y*-axis represents the logarithm of variance (log *V*), and the *x*-axis represents the logarithm of mean (logm). SPL is inherently accrual-based; both accrual variants — using accrual index (*i*) and cumulative sample count (*Q*) — are shown.

**figure 1B.**
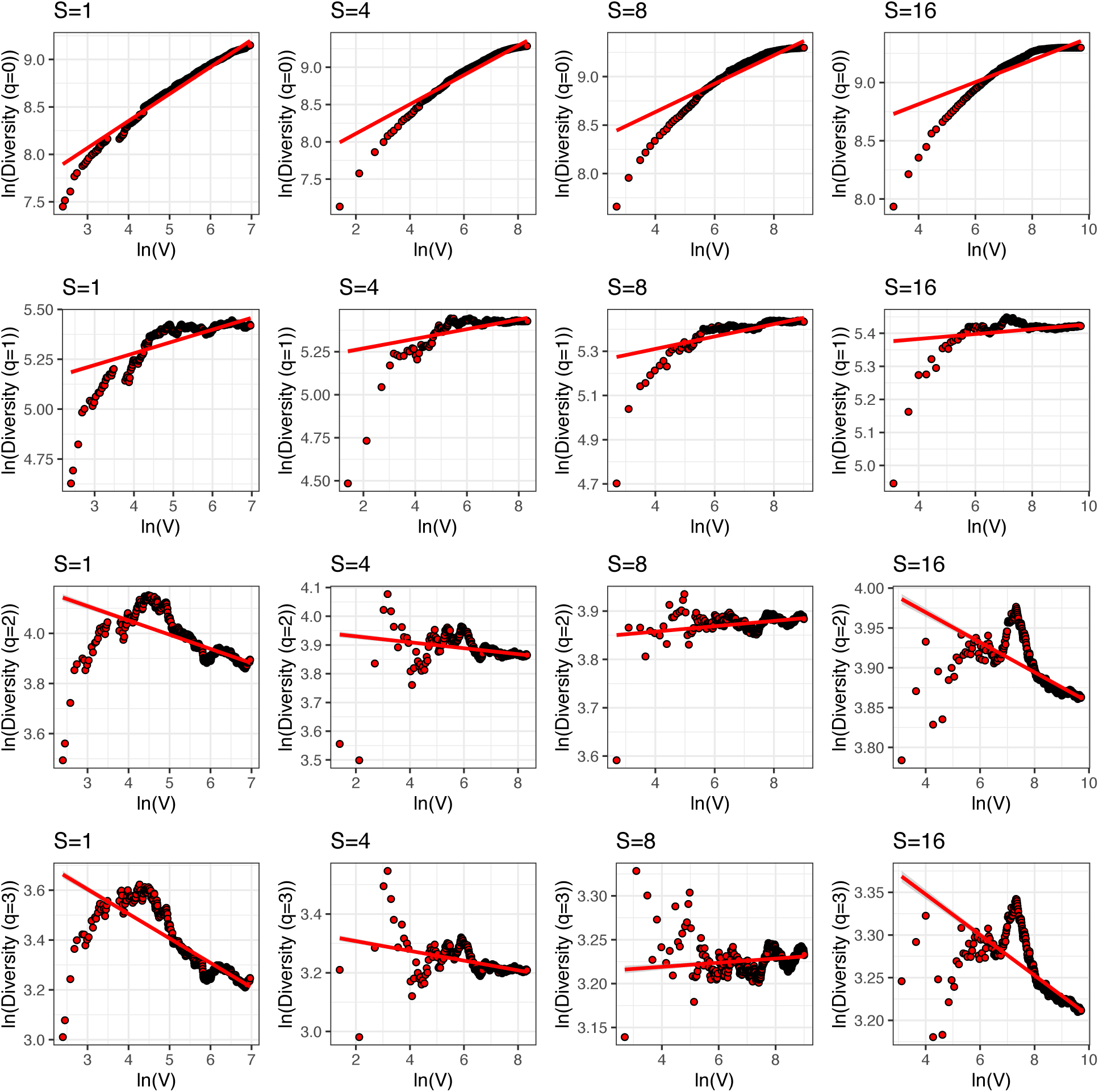
New DMR (diversity-mean relationship) models with sample accrual fitted to the AGP dataset at MUS scales *S* = 1,4,8,16. The y-axis represents the logarithm of Hill diversity [log *D*(*q*)] for orders *q* = 0,1,2,3, and the *x*-axis represents the logarithm of mean (*m*) of the accrued samples.

**figure 1C.**
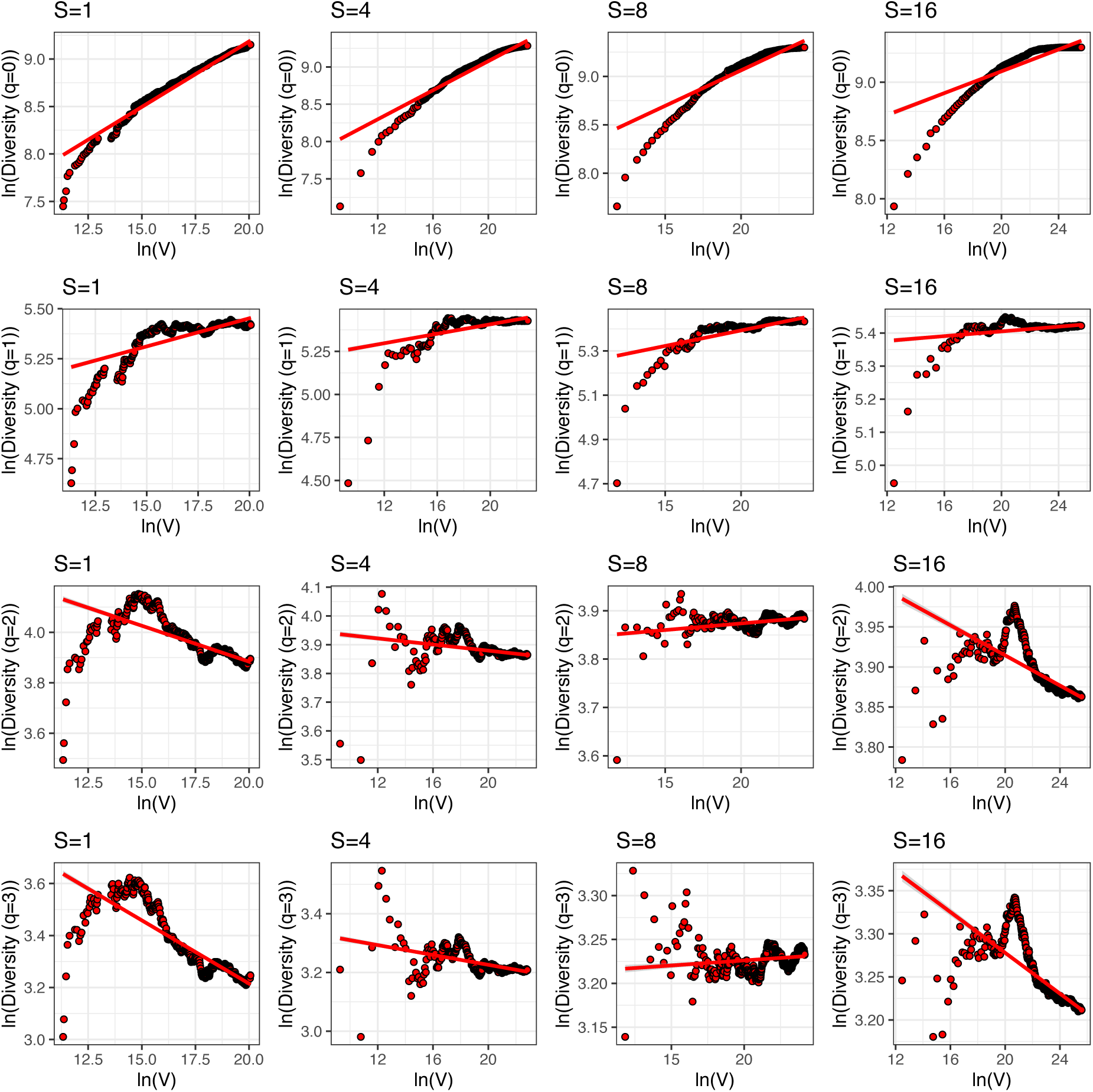
New DVR (diversity–variance relationship) models with sample accrual fitted to the AGP dataset at MUS scales *S* = 1,4,8,16. The y-axis represents the logarithm of Hill diversity [log *D*(*q*)] for orders *q* = 0,1,2,3, and the *x*-axis represents the logarithm of variance (logV) of the accrued samples.

**figure 1D.**
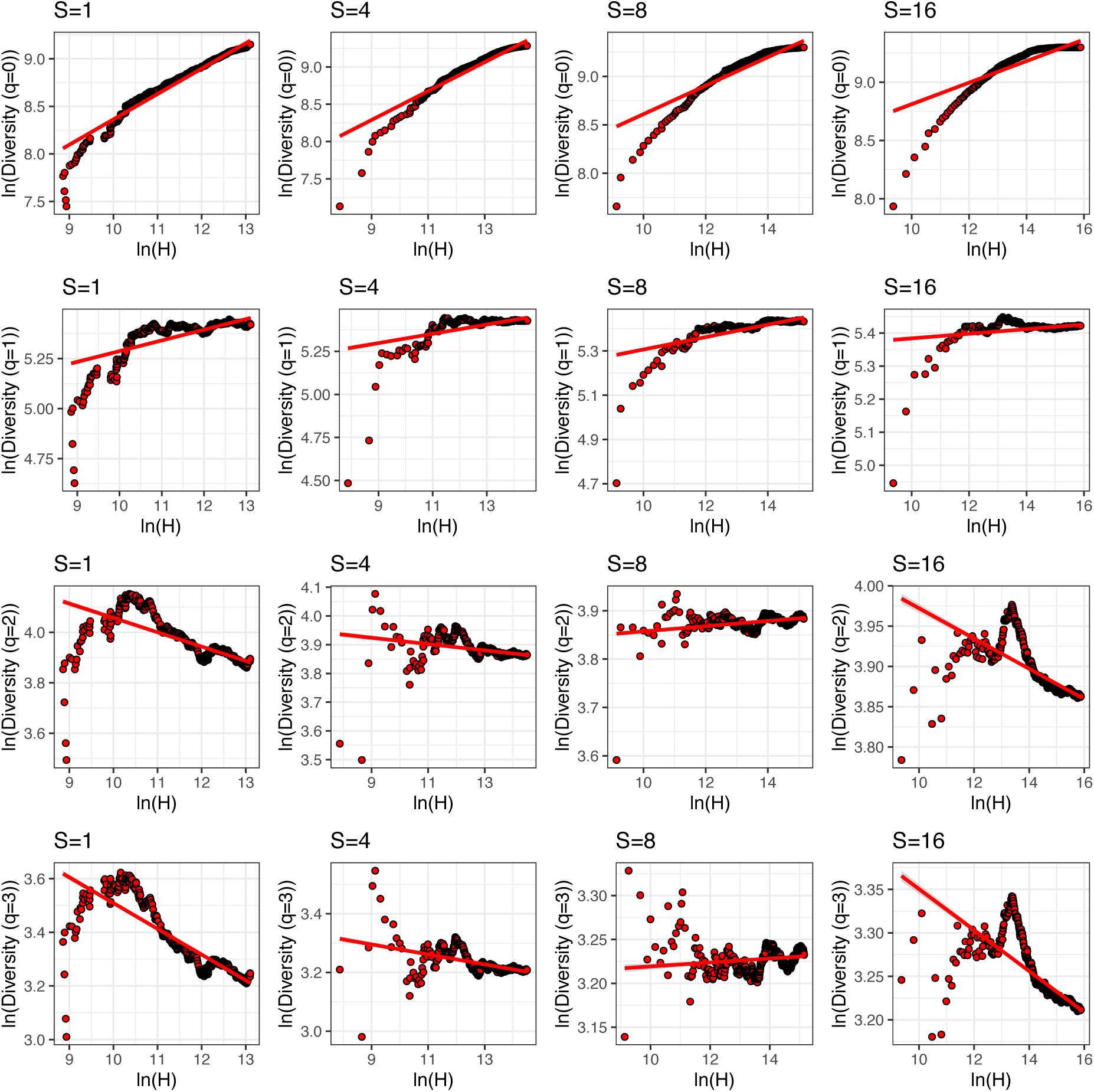
New DHR (diversity-heterogeneity relationship) models with sample accrual fitted to the AGP datasets at MUS scales *S* = 1,4,8,16. The y-axis represents the logarithm of Hill diversity [log *D*(*q*)] for orders *q* = 0,1,2,3, and the *x*-axis represents the logarithm of heterogeneity (V/m) of the accrued samples.

**figure 2A.**
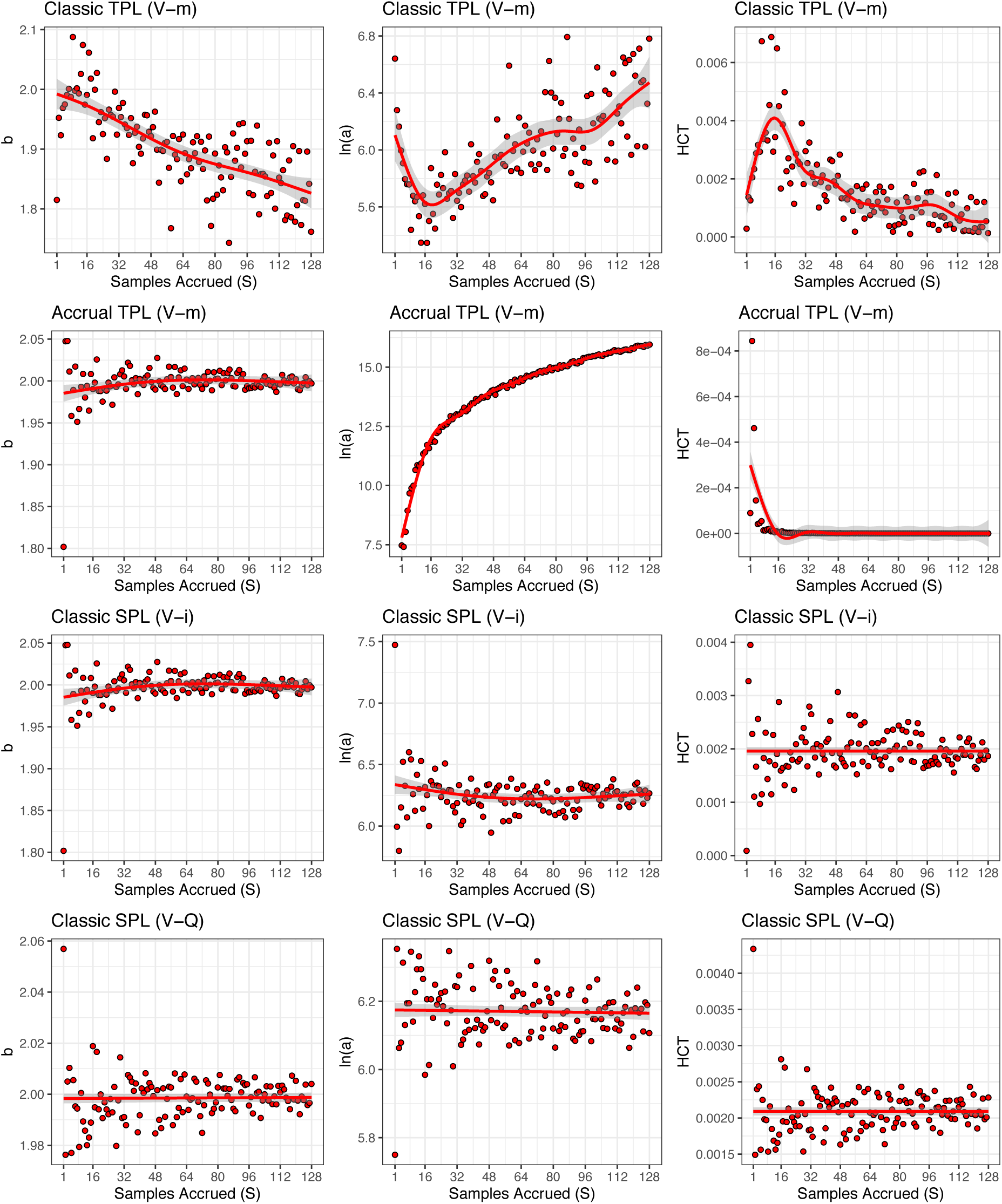
Relationships between the parameters (*b*, ln(*a*), HCT) of classic TPL, new accrual TPL, and classic SPL (*V* − *i* and *V* − *Q* variants) at multi-unit scale (MUS, *S* = 1 *to* 128) for the AGP dataset. The *x*-axis represents the MUS scale (S), and the y-axis represents the corresponding parameter values at each scale.

**figure 2B.**
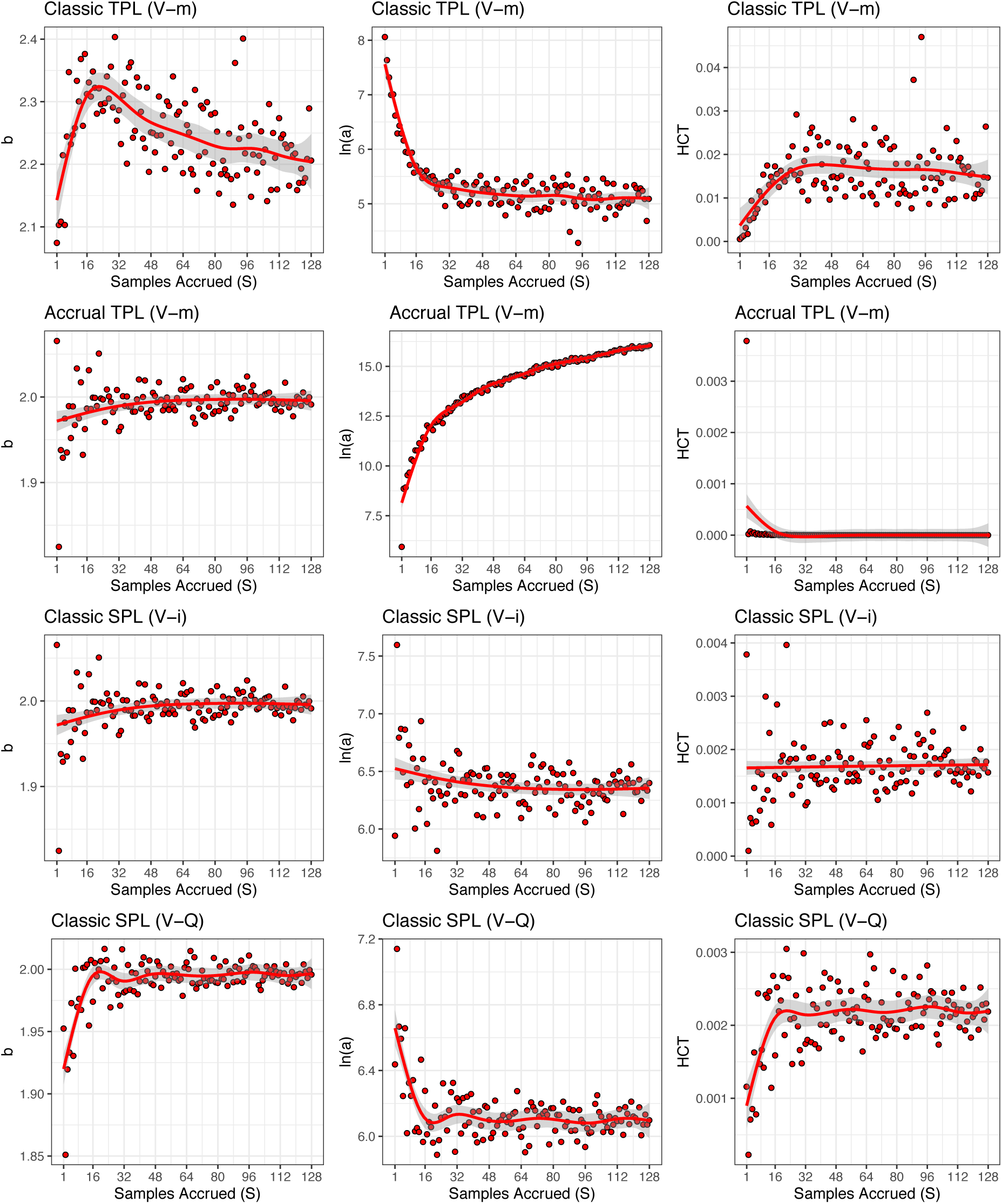
Relationships between the parameters (*b*, ln(*a*), HCT) of classic TPL, new accrual TPL, and classic SPL (*V* − *i* and *V* − *Q* variants) at multi-unit scale (MUS, *S* = 1 *to* 128) for the vaginal microbiome dataset. The *x*-axis represents the MUS scale (S), and the y-axis represents the corresponding parameter values at each scale.

**figure 3A.**
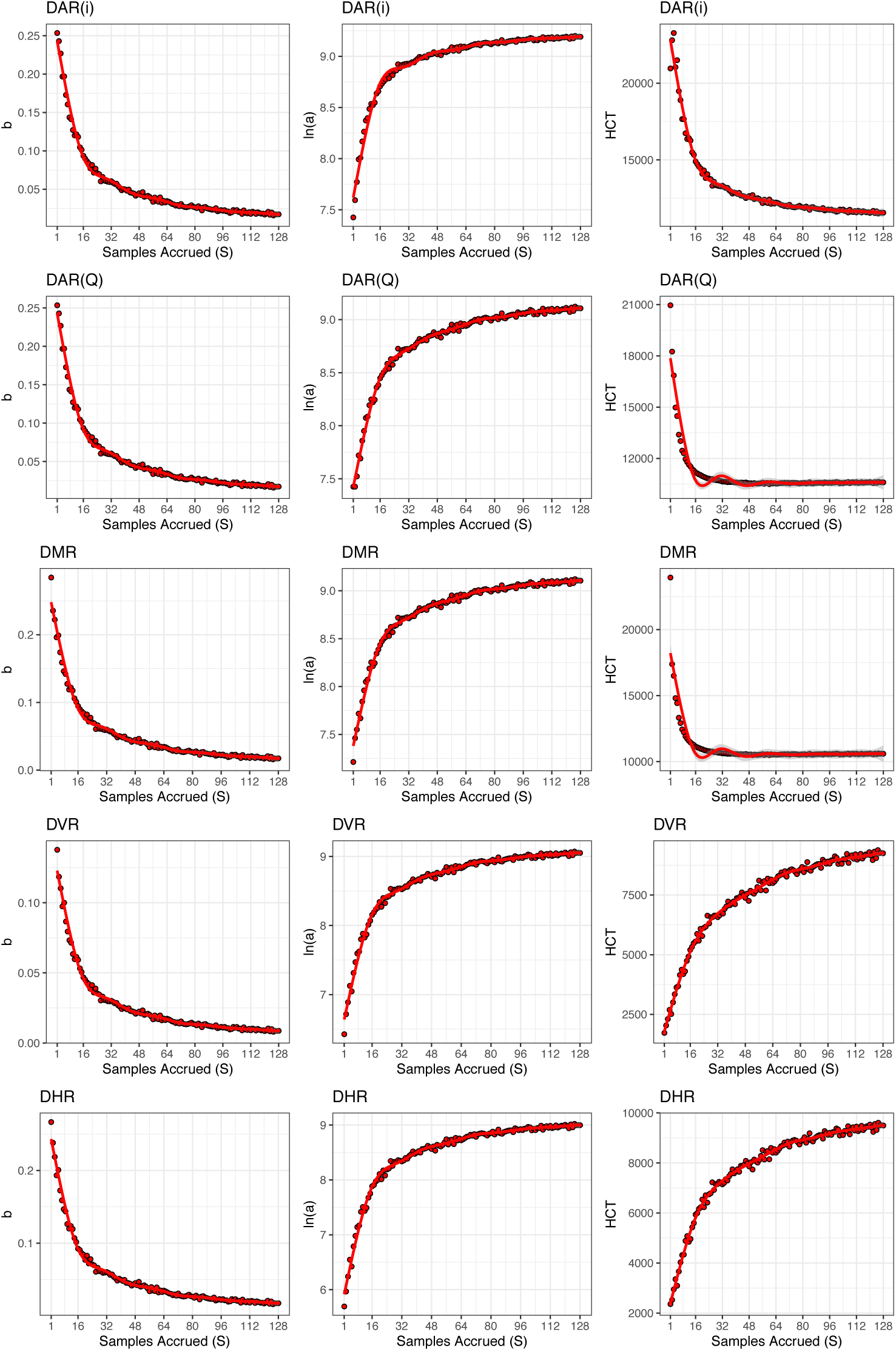
Relationships between the parameters [ *b*, log(*a*), *HCT*] of DAR (Diversity–Area Relationship), DMR (Diversity–Mean Relationship), DVR (Diversity–Variance Relationship), and DHR (Diversity–Heterogeneity Relationship) with diversity order (A = B) at multi-unit scales (MUS; *S*=1 to 128) for the AGP dataset. The *x*-axis represents the MUS scale (*S*), and the *y*-axis represents the corresponding parameter values at each scale.

**figure 3B.**
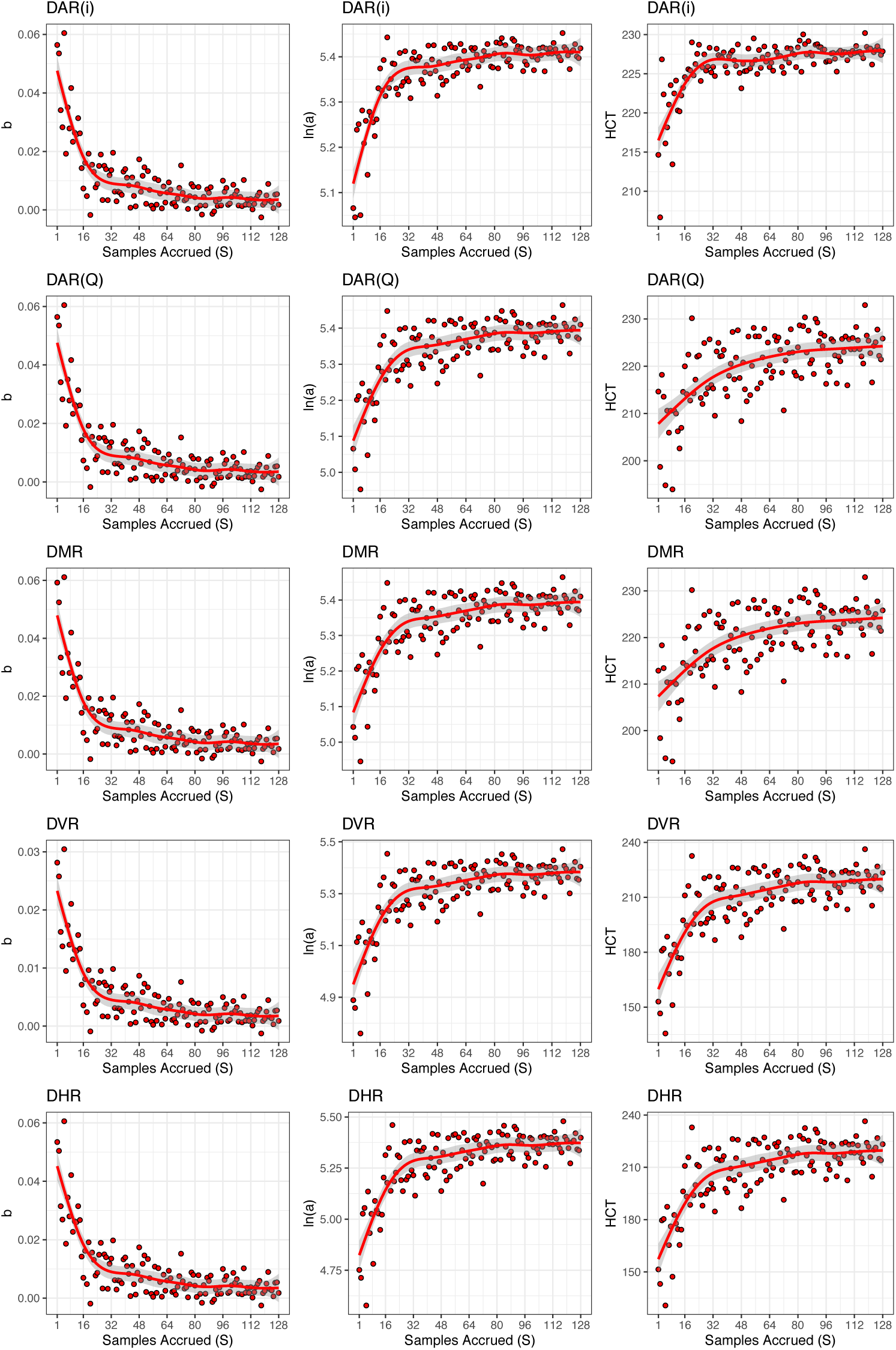
Relationships between the parameters [ *b*, log(*a*), *HCT*] of DAR (Diversity–Area Relationship), DMR (Diversity–Mean Relationship), DVR (Diversity–Variance Relationship), and DHR (Diversity–Heterogeneity Relationship) with diversity order (A = 1) at multi-unit scales (MUS; *S*=1 to 128) for the AGP dataset. The *x*-axis represents the MUS scale (*S*), and the *y*-axis represents the corresponding parameter values at each scale.

**figure 3C.**
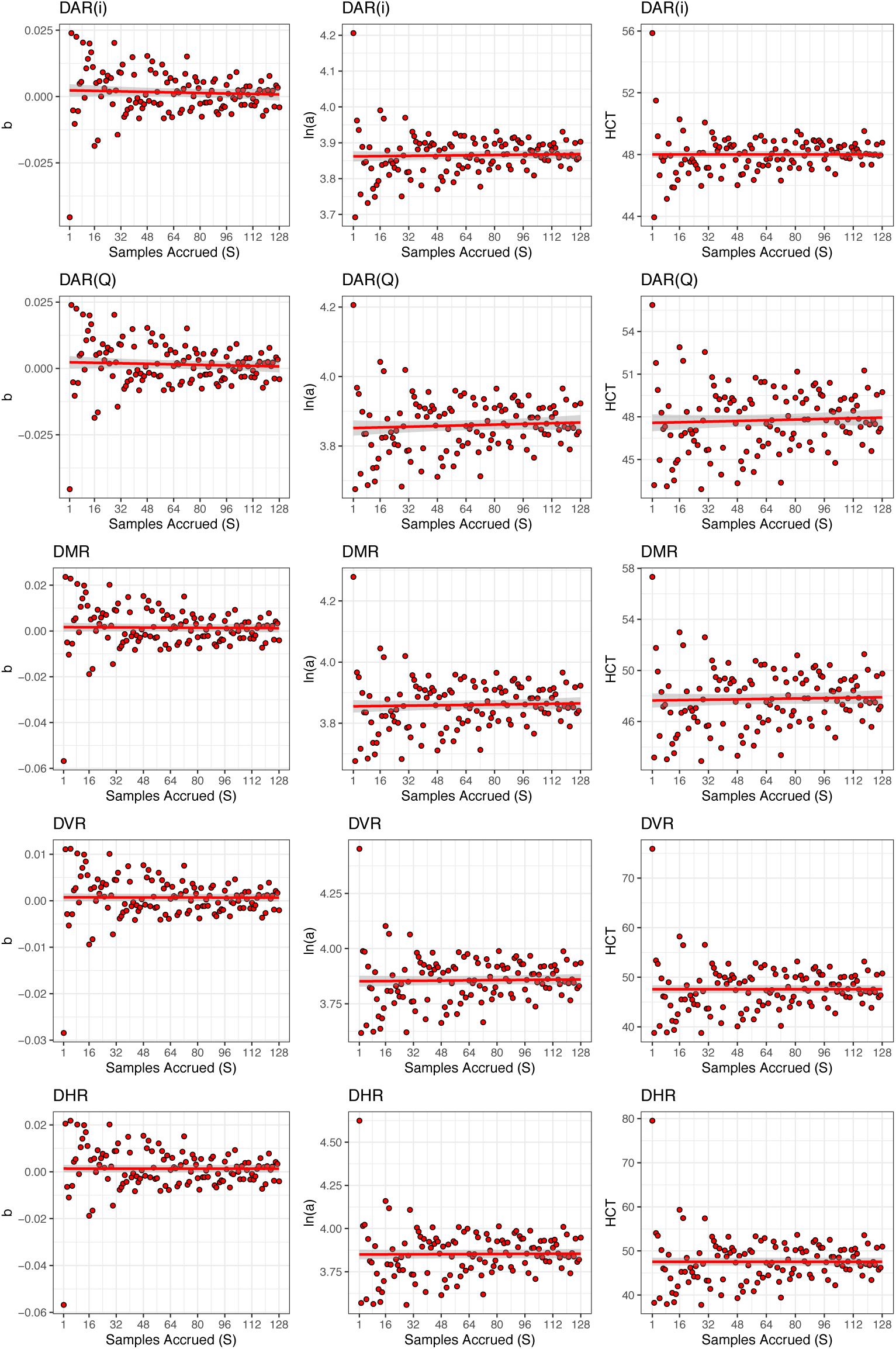
Relationships between the parameters [ *b*, log(*a*), *HCT*] of DAR (Diversity–Area Relationship), DMR (Diversity–Mean Relationship), DVR (Diversity–Variance Relationship), and DHR (Diversity–Heterogeneity Relationship) with diversity order (A = D) at multi-unit scales (MUS; *S*=1 to 128) for the AGP dataset. The *x*-axis represents the MUS scale (*S*), and the *y*-axis represents the corresponding parameter values at each scale.

**figure 3D.**
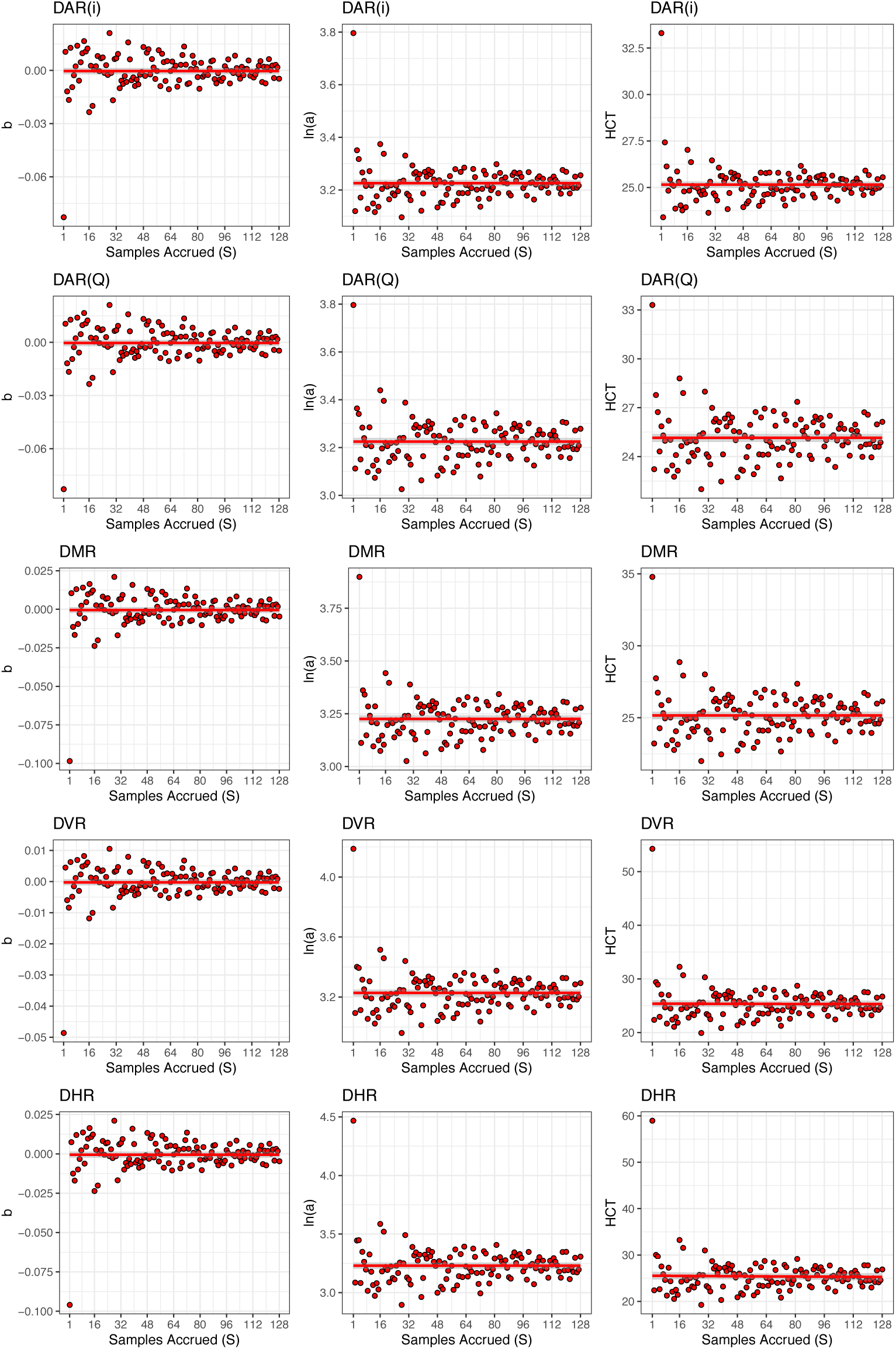
Relationships between the parameters [ *b*, log(*a*), *HCT*] of DAR (Diversity–Area Relationship), DMR (Diversity–Mean Relationship), DVR (Diversity–Variance Relationship), and DHR (Diversity–Heterogeneity Relationship) with diversity order (A = E) at multi-unit scales (MUS; *S*=1 to 128) for the AGP dataset. The *x*-axis represents the MUS scale (*S*), and the *y*-axis represents the corresponding parameter values at each scale.

**figure 4A.**
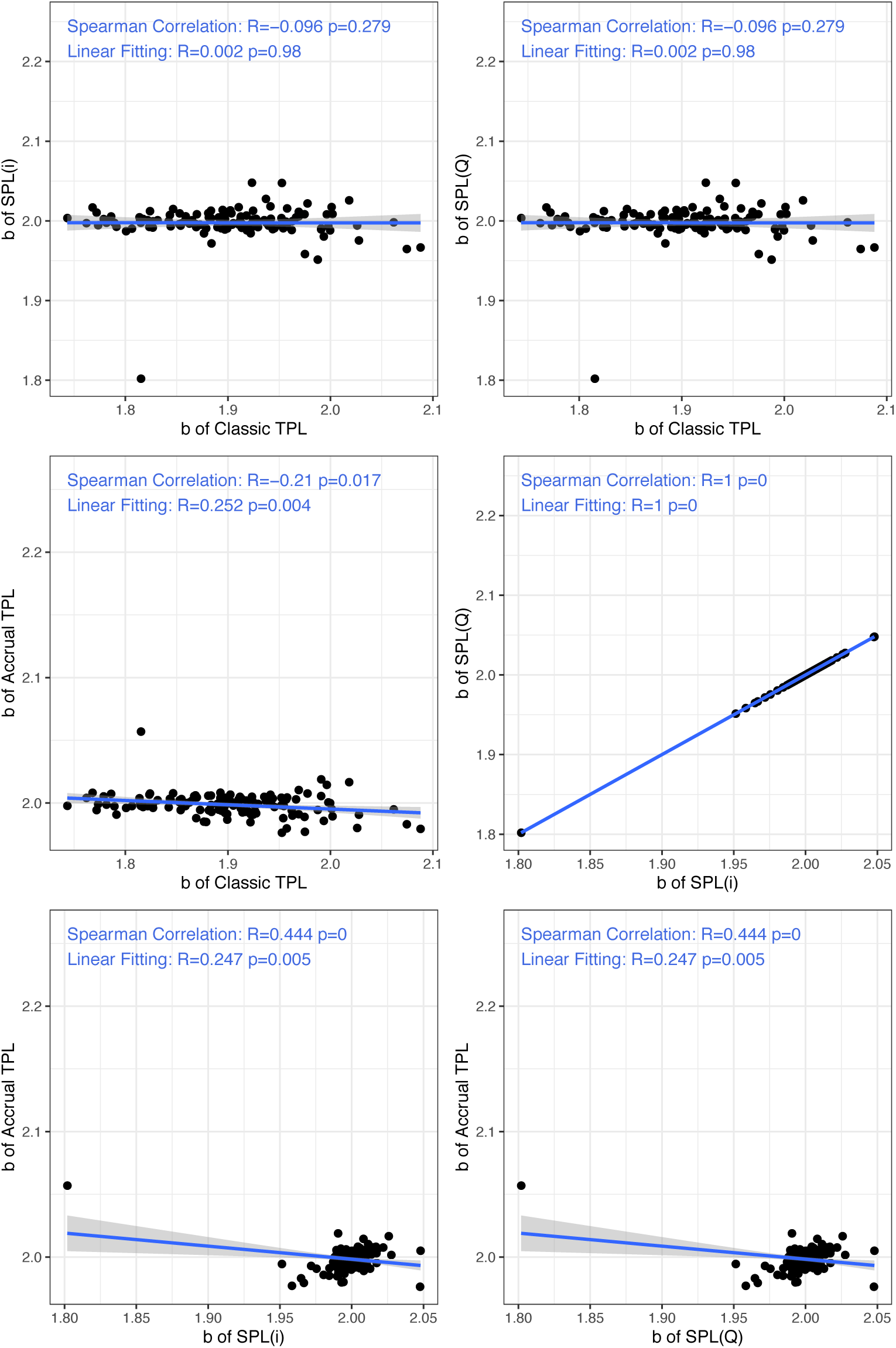
Spearman correlations of *b* values across accrual sample scales (MUS; S=1 to 128) between classic TPL, accrual TPL, SPL (V-i), and SPL (V-Q) models for the AGP dataset

**figure 4B.**
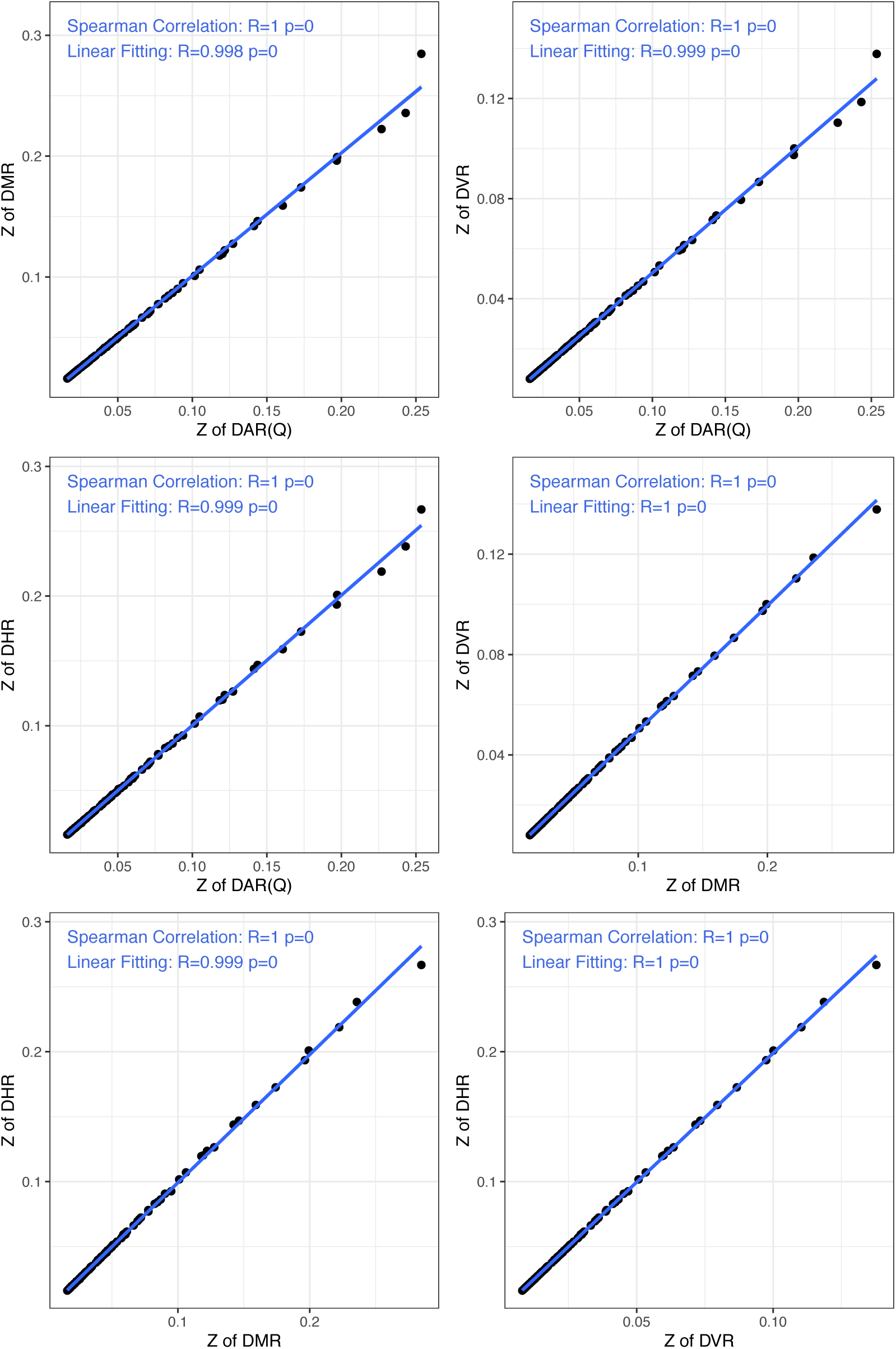
Spearman correlations of ***z*** value across accrual sample scales (MUS; S=1 to 128) between DAR, DMR, DVR, DHR models for the AGP dataset with diversity order *q*=0.

**figure 4C.**
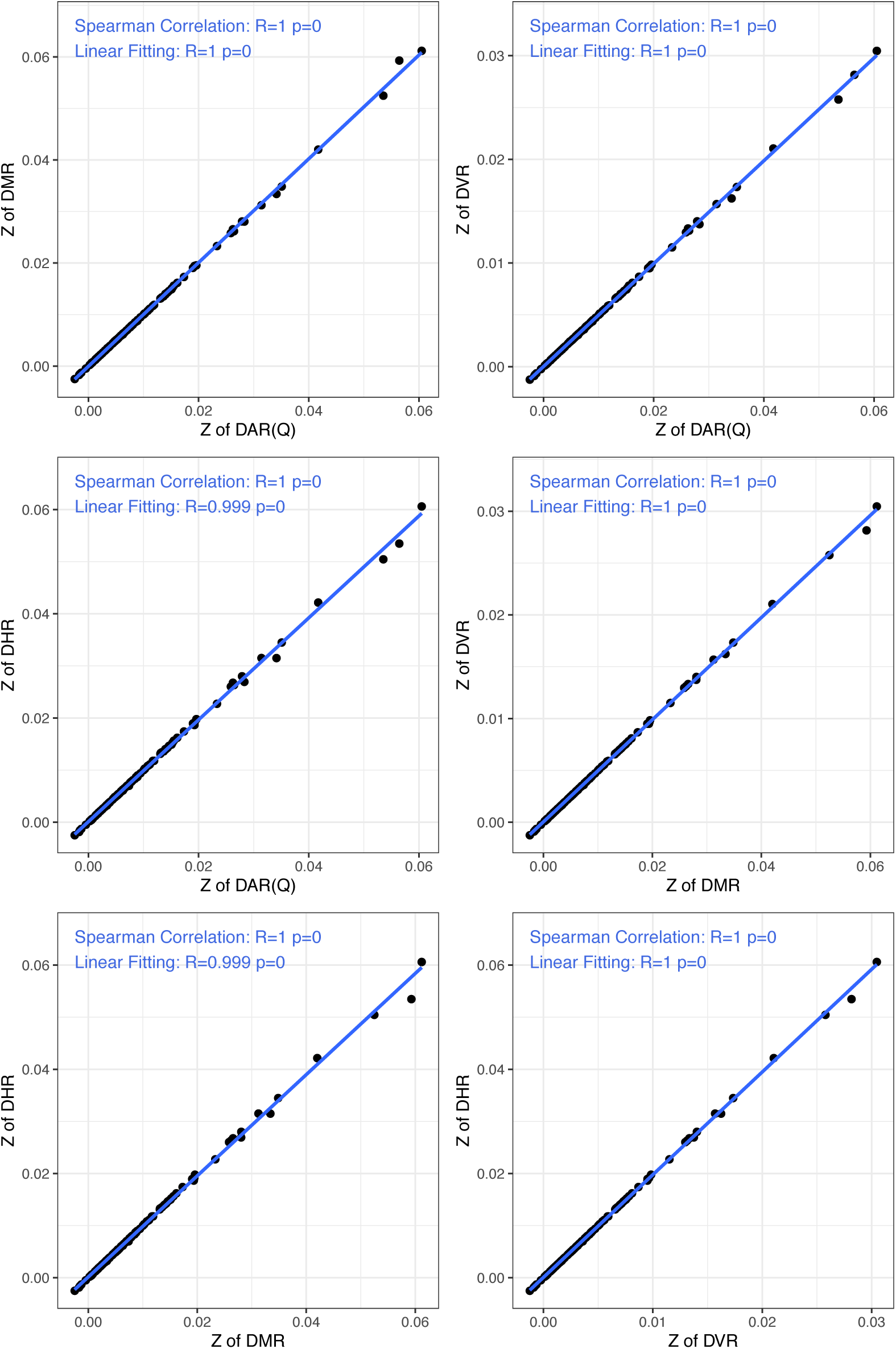
Spearman correlations of ***z*** value across accrual sample scales (MUS; S=1 to 128) between DAR, DMR, DVR, DHR models for the AGP dataset with diversity order *q*=1.

**figure 4D.**
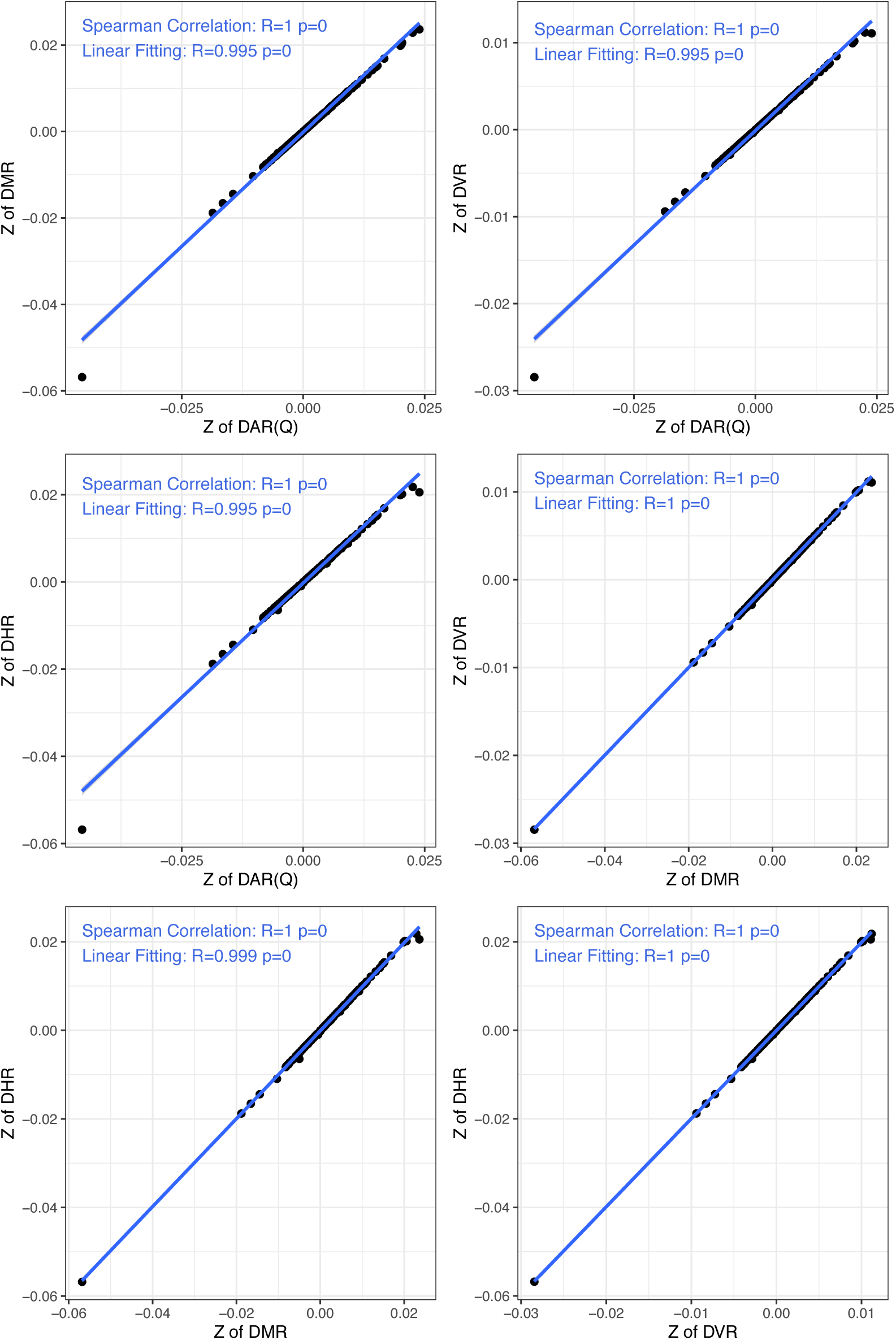
Spearman correlations of ***z*** value across accrual sample scales (MUS; S=1 to 128) between DAR, DMR, DVR, DHR models for the AGP dataset with diversity order *q*=2.

**figure 4E.**
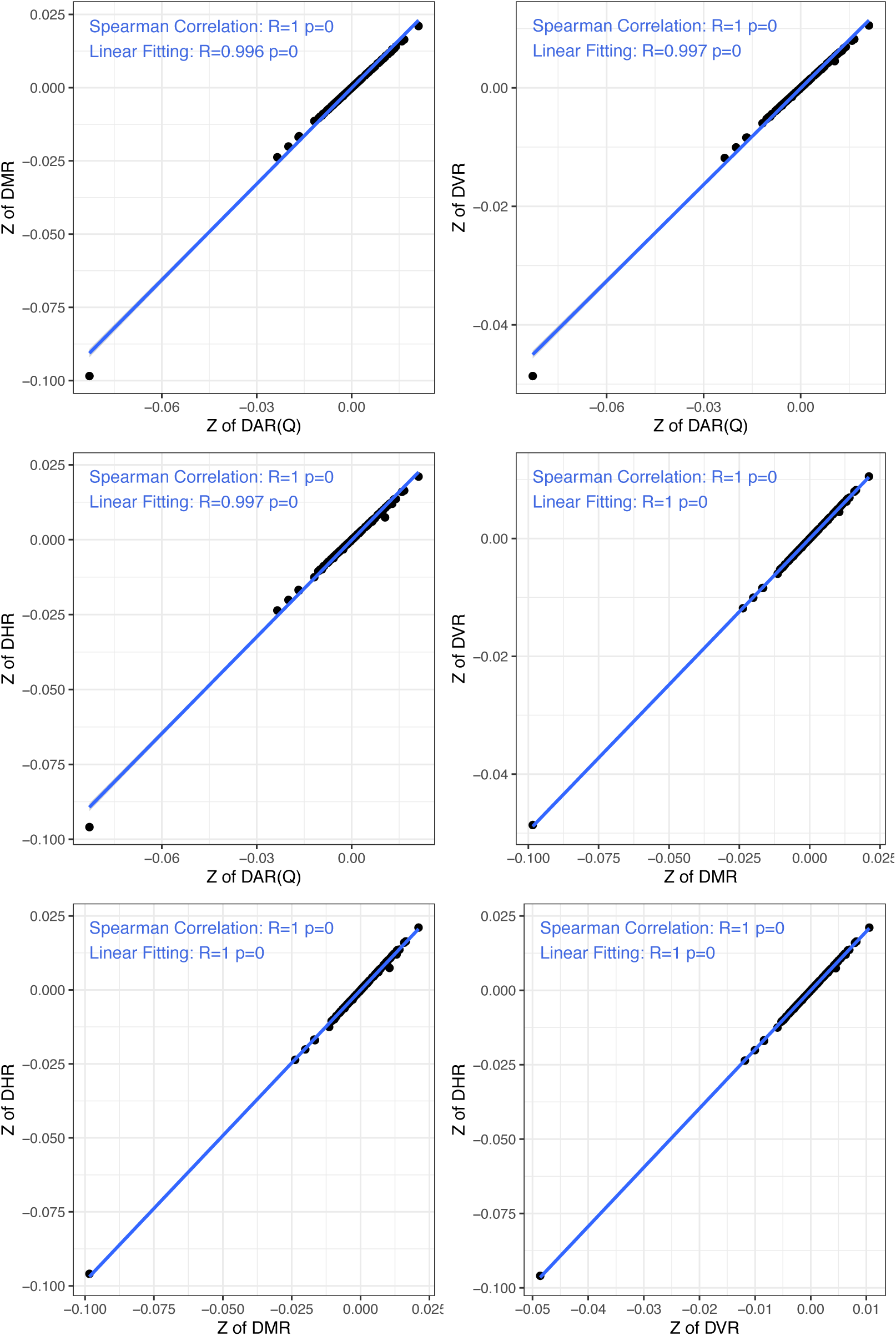
Spearman correlations of ***z*** value across accrual sample scales (MUS; S=1 to 128) between DAR, DMR, DVR, DHR models for the AGP dataset with diversity order *q*=3.

**figure 4F.**
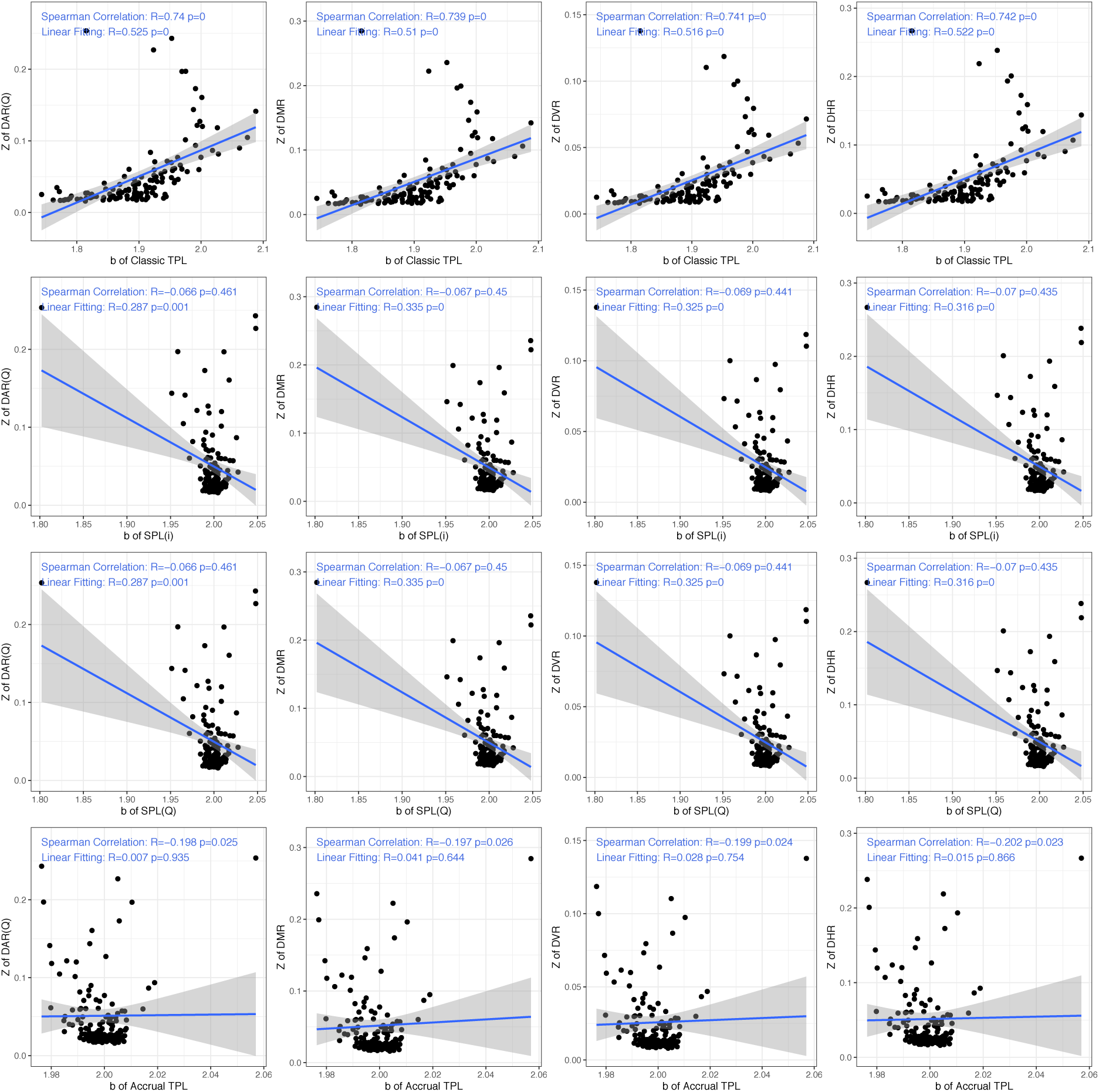
Spearman correlations between *b* values from classic TPL, accrual TPL, SPL (*i*), SPL (*Q*) models and the *z* values from DAR, DMR, DVR, DHR models across accrual sample scales (MUS; S=1 to 128) for the AGP dataset with diversity order *q*=0.

**figure 4G.**
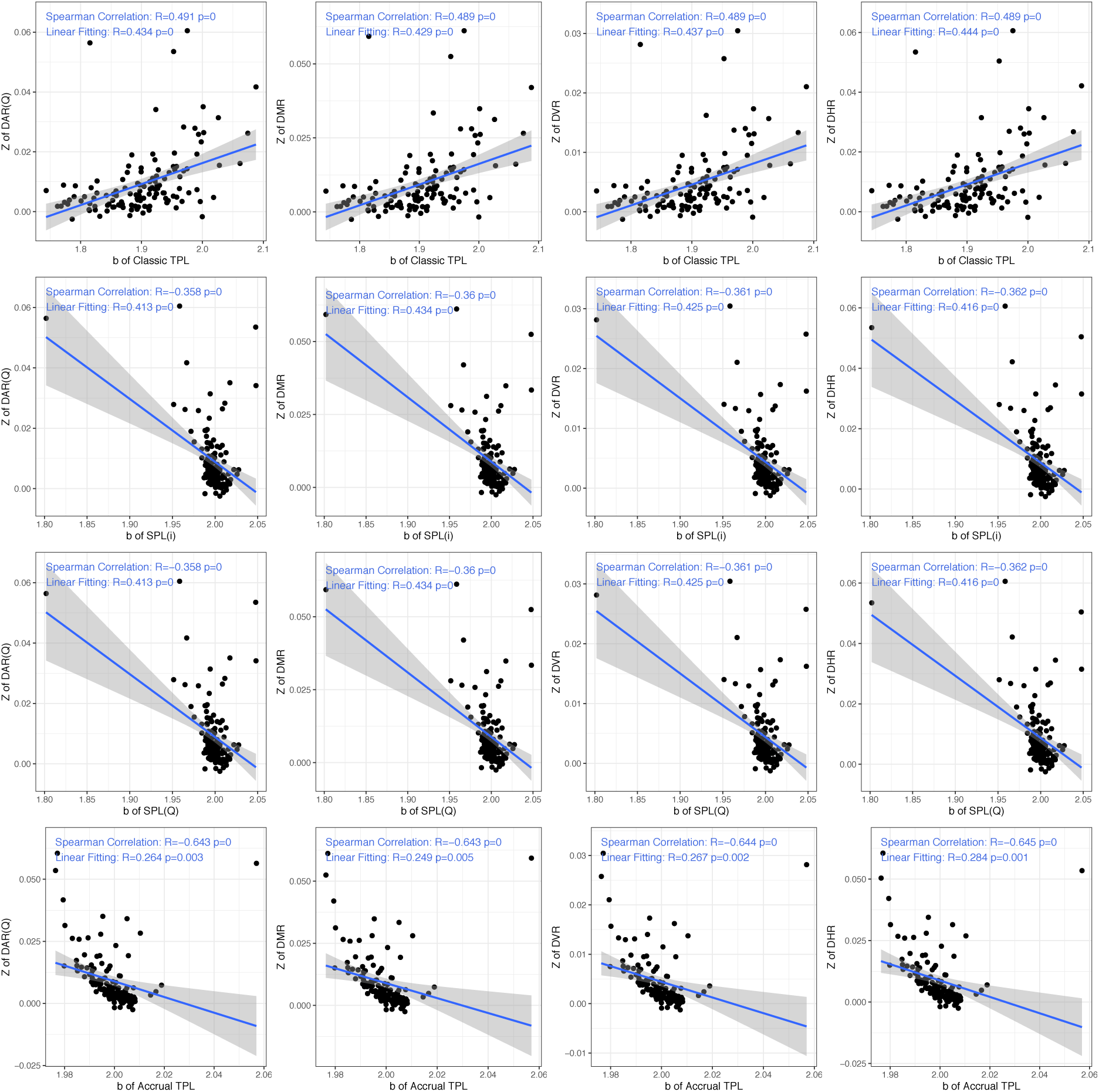
Spearman correlations between *b* values from classic TPL, accrual TPL, SPL (*i*), SPL (*Q*) models and the *z* values from DAR, DMR, DVR, DHR models across accrual sample scales (MUS; S=1 to 128) for the AGP dataset with diversity order *q*=1.

**figure 4H.**
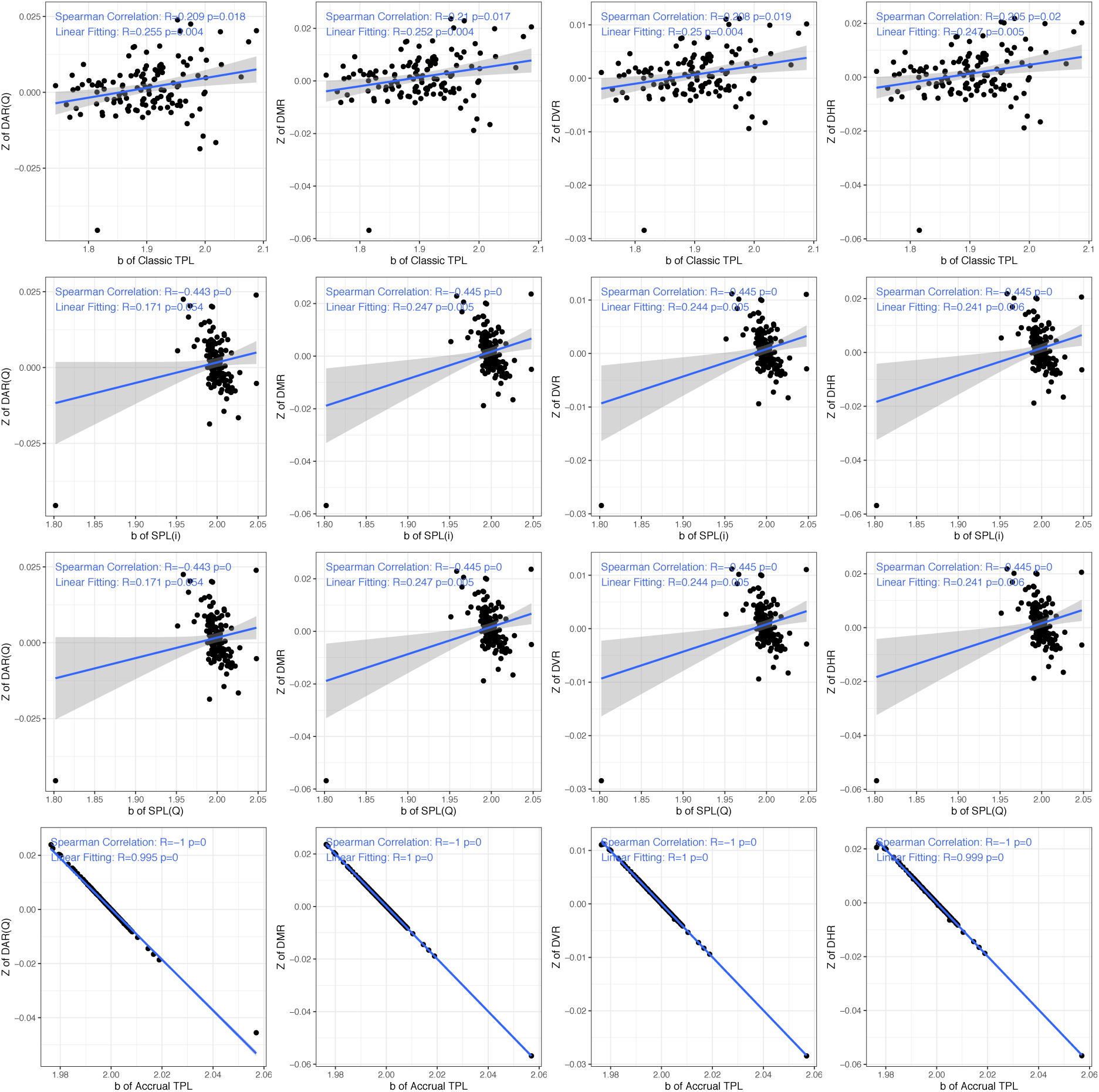
Spearman correlations between *b* values from classic TPL, accrual TPL, SPL (*i*), SPL (*Q*) models and the *z* values from DAR, DMR, DVR, DHR models across accrual sample scales (MUS; S=1 to 128) for the AGP dataset with diversity order *q*=2.

**figure 4I.**
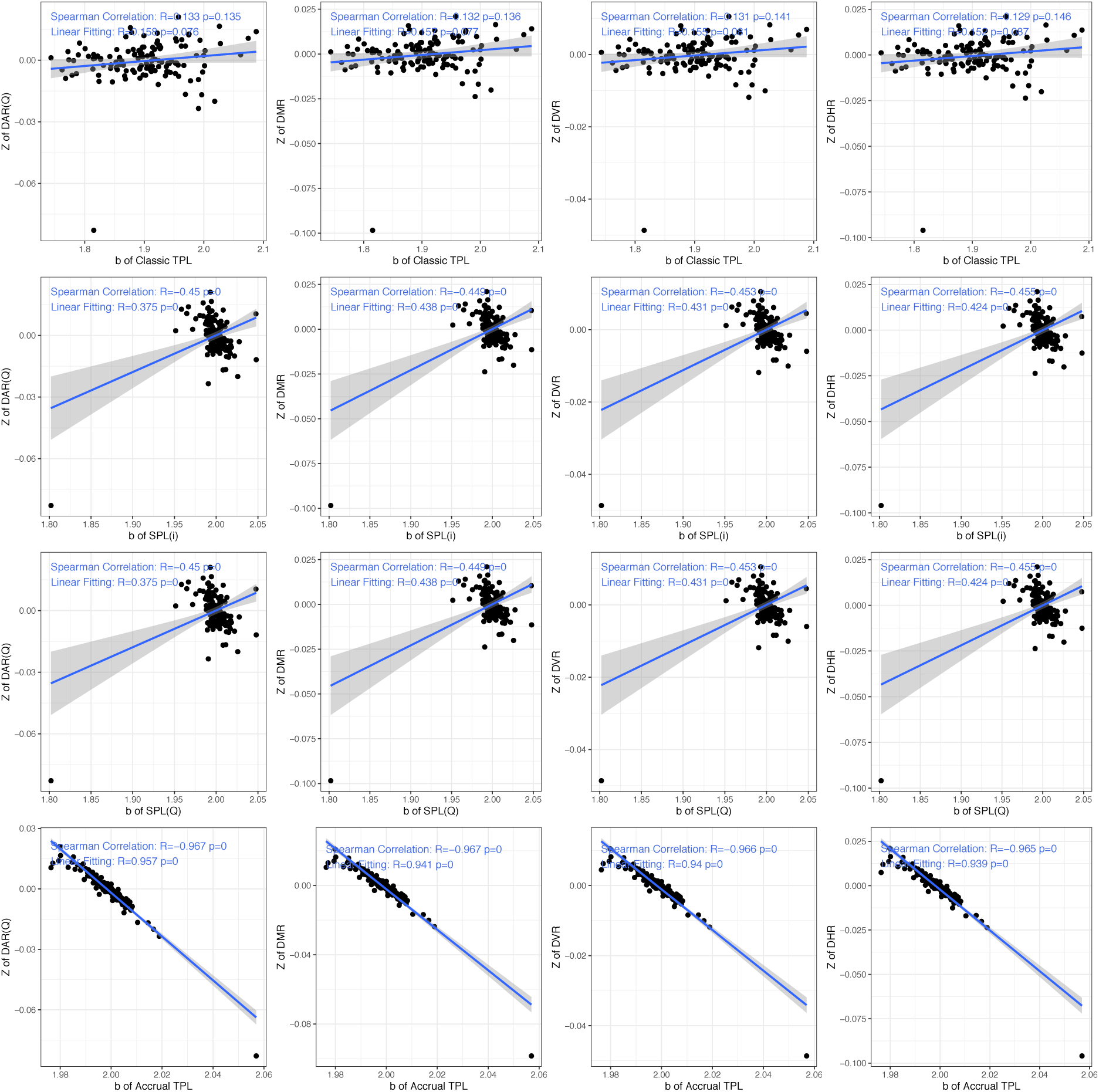
Spearman correlations between *b* values from classic TPL, accrual TPL, SPL (*i*), SPL (*Q*) models and the *z* values from DAR, DMR, DVR, DHR models across accrual sample scales (MUS; S=1 to 128) for the AGP dataset with diversity order *q*=3.

**Figure 5.**
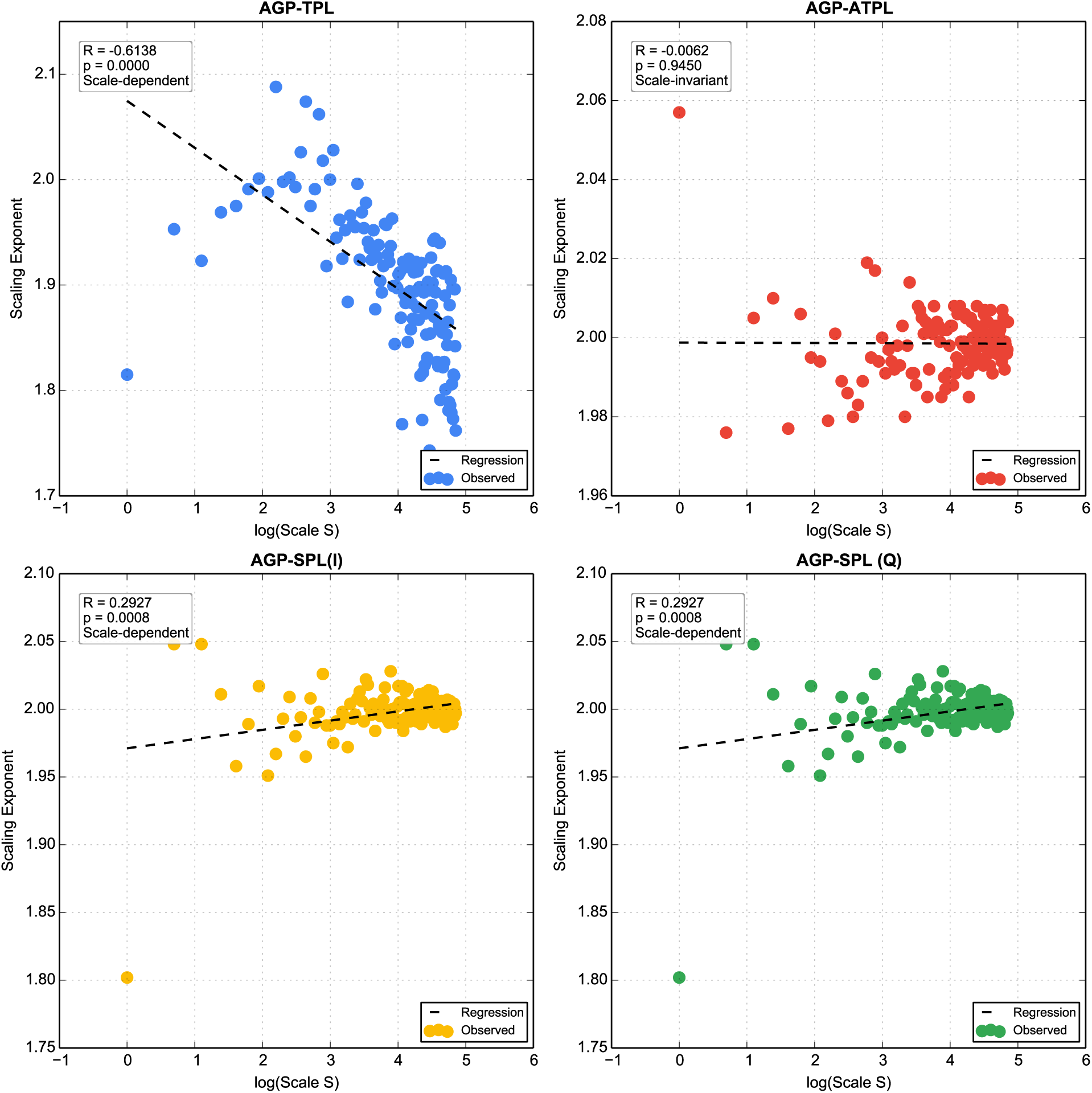
Scale invariance testing for AGP dataset with replacement sampling. Each panel shows the scaling exponent (*b*) plotted against log-transformed MUS scale (*S*) for four power law models: (A) classic TPL, (B) new accrual TPL, (C) SPL (V-i), and (D) SPL (V-Q). Solid circles represent observed values; dashed lines indicate linear regression fits. *R* and *p*-values are shown in each panel. Only accrual TPL (B) exhibited strict scale invariance (*p* > 0.05, slope not significantly different from zero), while the other three models showed varying degrees of scale dependence.

### 3.0 US-Scale Power Laws: Patterns at the Unit Scale (Individual Level)

#### 3.0.1 Classic TPL and Ecological Heterogeneity

Classic TPL at US scale (Table 2A) revealed strong fits in both habitats (*p* = 0.000). The exponent *b* was 1.815 (AGP) and 2.074 (vaginal), both exceeding 1, indicating variance grows faster than the mean (Ma, 2015). Vaginal exhibited higher b, consistent with its low diversity and *Lactobacillus* dominance. Intercept ln(a) was also higher in vaginal (8.060 vs. 6.642), reflecting greater baseline heterogeneity. HCT values were near zero in both habitats.

#### 3.0.2. Accrual TPL and SPL With Sample Accrual

With accrual (Table 2A), TPL *b* shifted oppositely: increased for AGP (accrual TPL 2.057 vs. non-accrual or classic TPL 1.815) but decreased for vaginal (1.952 vs. 2.074), indicating habitat-specific accrual effects. Accrual TPL fits were near-perfect (*p* = 0.000).

SPL yielded *b* = 1.802 (AGP) and 2.065 (vaginal), with excellent fits. At US scale, the two SPL variants (V-Index and V-Q) are identical. SPL intercepts differed from accrual TPL in both habitats. The similarity between SPL and accrual TPL *b* suggests both capture heterogeneity accumulation.

In summary, the three heterogeneity models at US scale reveal complementary facets: classic TPL captures cross-sectional spatial heterogeneity; accrual TPL captures cumulative heterogeneity; SPL captures environmental heterogeneity scaling. The direction of *b*-shift differs between habitats, foreshadowing classic TPL’s habitat-dependence and accrual-based models’ habitat-invariance at MUS scales.

#### 3.0.3. Diversity Power Laws: DAR, DMR, DVR, DHR

All four DPL models were fitted at US scale with accrual (Table 3A). Fits were excellent for *q* = 0,1 in both habitats. At higher orders, AGP fits remained excellent, while vaginal weakened: DVR at *q* = 3 was marginal (*p* = 0.026), and DHR at *q* = 3 was non-significant (*p* = 0.857).

For *q* = 0, DAR z was 0.254 (AGP) and 0.757 (vaginal). For *q* = 1, *z* declined to 0.056 and 0.135. AGP showed sign reversal at higher orders: *z* became negative at *q* = 2, 3, while vaginal remained positive but declining. This reflects Hill numbers’ sensitivity to rare vs. dominant species.

DMR, DVR, and DHR showed the same sign patterns: positive at low *q*, negative at high *q* for AGP; consistently positive for vaginal. At *q* = 0, DMR and DVR diverged (0.285 *vs*. 0.138 for AGP; 0.716 *vs*. 0.365 for vaginal). DHR was intermediate for AGP (0.267) but matched DAR for vaginal (0.737).

At US scale, *m*, *V*, and *V/M* are nearly perfectly correlated with diversity (Table S5, *S* = 1; R ≈ 0.9999 for *q* = 0). The divergence in exponents between DMR (0.285) and DVR (0.138) for AGP reflects the functional relationship *V* = *aM^b^*, not independent information. For vaginal, similar near-perfect correlations yield diverging exponents (0.716 vs. 0.365), consistent with its higher TPL exponent (*b* = 2.074).

### 3.1. Scale Dependence: MUS Power Laws Across S=1 to 128

#### 3.1.1. TPL Scale Invariance (or Lack Thereof)

To examine whether TPL is scale-invariant, we compared classic TPL (without accrual) and accrual TPL across MUS scales *S* = 1 to 128 (Table S1A). Standard errors were small (AGP classic TPL: *SE* ≈ 0.006; vaginal classic TPL: *SE* ≈ 0.006; accrual TPL: *SE* ≈ 0.001– 0.002), suggesting that observed variation reflects real differences rather than sampling error.

Classic TPL showed some evidence of scale-dependence. In AGP, *b* ranged from 1.815 (*S* = 1) to 2.088 (S=9S=9), with a mean of 1.902. In vaginal, b ranged from 2.074 (*S* = 1) to 2.404 (*S* = 30), with a mean of 2.248. The variation in *b* across *S*, including a pronounced peak in vaginal around *S* = 30, suggests that classic TPL may not be scale-invariant.

In contrast, accrual TPL for the AGP dataset showed near-constant *b* across all SS: mean *b* = 1.999 (range 1.976–2.057). The standard errors were considerably smaller than those for classic TPL, suggesting that accrual TPL may be scale-invariant — the rate of heterogeneity accumulation appears not to depend on grain size. Classic TPL, as traditionally applied, appears scale-dependent, while accrual TPL reveals a potentially scale-invariant property.

The preceding observations, based on standard error comparisons, are informal and should be interpreted with caution. Rigorous scale-invariance testing, using regression of the scaling exponent against log-transformed scale, revealed that fewer than 20% (approximately 17%) of the models examined were scale-invariant. Figure 5 illustrates four representative examples; among these, only accrual TPL for the AGP dataset demonstrated strict scale-invariance, consistent with earlier informal observations.

#### 3.1.2. SPL and Environmental Heterogeneity Scaling

To assess whether environmental heterogeneity scaling depends on grain (scale) size, we fitted SPL at MUS scale with sample accrual, using both accrual variants (V-Index and V-Q). The SPL exponent *b* was remarkably stable across *S*=1 to 128 in both habitats (Table S2A). For AGP, mean *b*≈1.998; for vaginal, mean *b*≈1.992. The near-constancy of *b* across scales indicates that environmental heterogeneity scaling is scale-invariant.

The two SPL accrual variants were mathematically equivalent, with identical *b* values and related intercepts: by ln(*a_v-q_*) = ln(*a_v-i_*) − *b* ⋅ ln (*S*), where *S* is the MUS size, *i.e.,* the grain size. The difference in ln(*a*) increased with SS, consistent with theory. The SPL exponent remained close to 2.0 in both habitats, contrasting with classic TPL but aligning with accrual TPL. The intercept for V-Index increased with *S*, reflecting increasing baseline variance. HCT remained near zero across all scales.

#### 3.1.3. DAR and Diversity Scaling Across Scales

To test whether diversity-area scaling depends on grain size, we fitted DAR at MUS scale with sample accrual across *S* = 1 to 128, for *q* = 0,1,2,3 (Table S3A). SAR is a special case of DAR at *q* = 0.

For species richness (*q* = 0), AGP *z* was positive across all *S*, ranging from 0.254 (*S* = 1) to 0.017 (*S* = 128), mean 0.051. The exponent declined with increasing S, indicating slower species accumulation at coarser grains. Vaginal *z* ranged from 0.757 (*S* = 1) to 0.04 (*S* = 128), mean 0.132, consistently higher than AGP.

At higher orders (*q* = 2,3), AGP showed negative z across most *S*, consistent with US-scale sign reversal; the magnitude decreased at larger S. Vaginal z remained positive across all *S* but declined with increasing *S*. Under the DAR framework (Ma, 2018), the decline in z with increasing *q* reflects the transition from richness-based to dominance-based diversity. Fit quality was high: 100% for *q=0*; 94.5% for AGP and 92.9% for vaginal *at q=3*.

#### 3.1.4. DMR, DVR, DHR at Multiple Unit Scales (MUS)

We next examined the three newly proposed DPL models — DMR, DVR, and DHR — at MUS scale with sample accrual across *S* = 1 to 128, for *q* = 0,1,2,3 (Table S4A). Mirroring US-scale results, all three models exhibited the same qualitative sign patterns: AGP positive at *q* = 0,1, negative at *q* = 2,3; vaginal positive across all *q*, declining with increasing *q*. This habitat contrast persisted across all MUS scales.

Magnitudes varied systematically with *S* (Table S4A), though signs remained stable. For AGP at q=0, DMR decreased from 0.285 to ∼0.018 with an average 0.0513 and standard error of 0.0043; DVR from 0.138 to ∼0.009; DHR from 0.267 to ∼0.017. For vaginal, similar declines were observed. DMR and DVR diverged (*e.g.,* AGP *q=0*: 0.285 vs. 0.138), reflecting *V* = *aM^b^*. DHR was intermediate for AGP but matched DAR for vaginal. Fit quality remained high (94.5–100% for AGP; 92.2–100% for vaginal).

Table S5 summary statistics across MUS scales reinforce these findings. For AGP, 100% of correlations between m, V, V/M, and diversity were significant. Positive correlations dominated at *q=0* (100%) but shifted sharply at higher orders: at *q* = 3, 44.5% were negative, consistent with the sign reversal in DPL exponents. For vaginal, significant correlations remained high (92–100% across *q*), but positive correlations declined from 100% at q=0 to 43.8% at q=3, reflecting weakening but persistent positive diversity scaling.

In contrast with SPL results, DPL models, including DAR, DMR, DVR, and DHR, are clearly not scale-invariant.

### 3.2. Cross-Model Relationships: Linking Heterogeneity and Diversity

Having established the scaling properties of individual power law models, we now address the central question: **how are heterogeneity scaling and diversity scaling related?** Three questions guide this section. First, how do different heterogeneity models (classic TPL, accrual TPL, SPL) relate to each other? Second, are the four diversity models (DAR, DMR, DVR, DHR) redundant or unique? Third, does heterogeneity scaling predict diversity scaling, and does it depend on scale or diversity order q? We use Spearman rank correlations across MUS scales (Table S6), with US scale included as S=1.

#### 3.2.1. Classic TPL, Accrual TPL and SPL

To assess whether different approaches capture shared or distinct information, we calculated Spearman’s correlations between classic TPL, accrual TPL, and SPL (Table S6).

##### Exponent *b*

Classic TPL showed no significant correlation with any accrual-based model (all p>0.05), confirming that the newly proposed accrual TPL is not redundant with classic TPL. The two SPL variants (Index-V and Q-V) were perfectly correlated (R=1.000, p=0.000), confirming their mathematical equivalence in terms of *b*. SPL variants correlated moderately with accrual TPL (R=0.444–0.728, p=0.000), suggesting that accrual TPL and SPL share partial information — indicating that the relationship between ecological and environmental heterogeneity is cumulative or additive in nature.

##### Intercept ln(a)

Classic TPL showed a strong positive correlation with Index-V accrual in gut (R=0.678, p=0.000) but a negative correlation in vaginal (R=−0.592, p=0.000), revealing a striking habitat-dependent reversal across MUS scales. Classic TPL vs. accrual TPL showed weak or non-significant correlations. Q-V vs. m-V (accrual) showed moderate positive correlations in both datasets (R=0.471–0.733, p=0.000).

##### Heterogeneity critical threshold (HCT)

Patterns mirrored ln(a): classic TPL vs. Index-V was strongly correlated in gut (R=0.714, p=0.000) but not in vaginal; classic SPL and accrual TPL showed moderate positive correlations in both habitats (R=0.461–0.733, p=0.000).

##### Key insights

(i) The scaling exponent *b* from classic TPL does not predict accrual-based exponents, but SPL and accrual TPL exponents are correlated. This suggests that the relationship between ecological heterogeneity and environmental heterogeneity is cumulative or additive. If we assume that environmental heterogeneity is the cause and ignore the counter-influence of ecological heterogeneity on environmental heterogeneity, this finding predicts that ecological heterogeneity should not be scale-invariant due to additive or cumulative effects. (ii) Intercept parameter [ln(*a*)] are highly sensitive to both accrual method and host site, revealing fundamental differences in baseline heterogeneity between gut and vaginal microbiomes. (iii) HCT, Only the Q-V variant of SPL (variance vs. cumulative sample count) showed moderate to strong positive correlations with accrual TPL (R=0.471–0.733, p=0.000), indicating that cumulative sample count captures similar information to mean abundance under accrual. The Index-V variant showed no significant correlation with accrual TPL. This again confirms that the relationship between ecological and environmental heterogeneity is cumulative or additive, and justifies the adoption of both SPL variants.

#### 3.2.2. DPL Model Inter-Comparisons

To assess possible redundancy among the four DPL models (DAR, DMR, DVR, DHR), we compared their scaling exponents z at US scale (Table 3A) and MUS scale (Table S5A and S6).

##### US scale (Table 3A)

At US scale, for AGP at *q* = 0, scaling exponents (*z*) varied: DAR (0.254), DMR (0.285), DVR (0.138), and DHR (0.267). However, sign patterns were identical across all four models: positive at *q* = 0,1; negative at *q* = 2,3 for AGP; positive at all *q* for vaginal.

##### MUS scale (Table S6A)

At MUS scale, for AGP at *q* = 0, exponents of all four models were nearly perfectly correlated (R = 1.000, *p* = 0.000), indicating that at species richness level, the four models carry essentially identical information across grain sizes. At higher orders (*q* = 1,2,3), correlations remained high (mostly R > 0.99), with only slight deviations at q=3 for vaginal (*e.g.,* DAR vs. DHR: R = 0.990).

##### Scale comparison

The high correlations at MUS scale contrast with the divergence in exponent magnitudes at US scale (e.g., DMR *z* = 0.285 vs. DVR *z* = 0.138 for AGP at q=0). This suggests that while the four models capture different absolute scaling rates at a single scale, their relative behavior across scales is highly coordinated. The sign patterns — the fundamental qualitative contrast between habitats — were identical at both scales.

##### Implication

The high redundancy among DAR, DMR, DVR, and DHR at MUS scale suggests that researchers may not need to fit all four models when interested in cross-scale behavior. DAR, being the simplest and most directly interpretable, may suffice for many applications. However, DHR holds a special place: it uses *H* = *V*/*M*, the variance-to-mean ratio, a direct measure of heterogeneity, making it the most natural bridge between diversity scaling and the heterogeneity scaling captured by TPL and SPL. While DMR and DVR can reveal habitat-specific differences in how diversity responds to mean species abundance versus variance, DHR directly tests the diversity–heterogeneity nexus without the confounding effects of mean abundance. For studies focused on the relationship between diversity and heterogeneity, DHR may be the preferred model.

#### 3.2.3 Habitat Contrasts: Gut *vs*. Vaginal Microbiomes

Do scaling laws differ between the high-diversity gut microbiome and the low-diversity, dominance-driven vaginal microbiome? We address this using Wilcoxon rank-sum tests comparing parameter estimates between habitats (Table S7A).

##### Heterogeneity comparisons

All four heterogeneity models — classic TPL, accrual TPL, and the two SPL variants (SPL-*i* and SPL-*Q*) — showed significant differences in the scaling exponent *b* between AGP and vaginal microbiomes, with classic TPL exhibiting the strongest difference. For the SPL-*i* variant, ln(a) and HCT showed no significant differences between the two microbiome types, but its counterpart SPL-*Q* model showed significant differences again.

##### Diversity comparisons

Similar to heterogeneity comparisons, all DPL parameter comparisons showed significant differences between the two microbiome types, with the exception of ln(a) in one case.

### 3.3. Does Heterogeneity Scaling Predict Diversity Scaling?

To address the central question of the diversity–heterogeneity nexus, we correlated heterogeneity exponents (TPL *b*, SPL *b*) with diversity exponents (DPL *z*) at MUS scale (Table S6A), where multiple *S* values enabled correlation analysis.

#### Classic TPL vs. DPL

Classic TPL b was strongly positively correlated with DAR *z* at *q* = 0 (R≈0.74, *p* = 0.000), but correlations weakened at higher *q* (e.g., *q* = 2: R ≈ 0.21, *p* = 0.018). For vaginal, correlations were weaker overall (*q* = 0: R ≈ 0.43, *p* = 0.000; *q* = 2: non-significant, *p* = 0.206). This suggests that classic TPL predicts diversity scaling primarily for *species richness*, and more strongly in gut than vaginal.

#### Accrual TPL vs. DPL

For AGP at *q* = 1, accrual TPL b was strongly negatively correlated with DAR z (R ≈ −0.64, *p* = 0.000); at *q* = 2, the correlation was perfect negative (R = −1.000, *p* = 0.000) with all four DPL models from DAR to DHR. For vaginal, similar patterns were observed with slightly weaker magnitudes. These strong negative correlations suggest that as ecological heterogeneity (accrual TPL *b*) increases, diversity scaling exponents z decrease — a potentially general principle of the diversity–heterogeneity nexus.

#### SPL vs. DPL

SPL *b* showed moderate to strong negative correlations with DPL *z*, particularly at high *q* in vaginal, consistent with the reciprocity principle between environmental heterogeneity and ecological diversity and heterogeneity (Ma & Ellison, 2026).

#### Conclusion

The essential difference between classic and accrual TPL is one of scale: classic TPL operates at the **community scale** (heterogeneity among individual hosts in the case of human microbiomes), while accrual TPL operates at the **metacommunity scale** (heterogeneity accumulation). These distinct scales explain their opposite relationships with diversity scaling.

For classic TPL (community scale), higher b was positively associated with higher DPL *z* at *q* = 0, indicating that communities with greater local heterogeneity exhibit faster species accumulation. For *q* > 0, correlations weakened or became non-significant. This suggests that while greater local heterogeneity may promote total species richness, but the effects should decline for dominant (*q* = 2) and elite species (*q* = 3).

For accrual TPL (metacommunity scale), higher *b* was negatively correlated with DPL z for virtually all q, and the magnitude of correlation increases at higher *q*. This indicates that faster heterogeneity accumulation flattens diversity scaling — a stabilization effect — with the strongest effect on diversity measures that emphasize dominant species.

In summary, while TPL predicts the relationship between ecological heterogeneity and diversity, SPL predicts the relationships between environmental (habitat) heterogeneity and diversity. For SPL, moderate to strong negative correlations with DPL z were observed, particularly at high *q* in vaginal microbiome, consistent with the reciprocity principle (Ma & Ellison, 2026). Ma & Ellison (2026) proposed that ecological and environmental heterogeneity are reciprocal. Our results support this principle: heterogeneity scaling (captured by TPL and SPL) predicts diversity scaling (captured by DPL). The tight coupling at *q*=0 and the systematic sign reversal at higher *q* demonstrate that the diversity–heterogeneity nexus follows predictable power law relationships, with the sign and strength determined by diversity order and habitat. One particularly interesting observation is the statistical goodness-of-fit of DHR (diversity–heterogeneity relationship) models across diversity orders (*q*) and habitat types (AGP vs. vaginal microbiomes), which relates diversity in Hill numbers to ecological heterogeneity measured *V*/*M*.

### 3.4. Modeling Results from the Sampling Scheme with Replacement

As mentioned in Methods section, both sampling schemes (with and without replacement) have advantages and disadvantages. All previous expositions are based on the scheme with replacement, which offers the advantage of a large sample size, specifically allowing us to investigate MUS from 1 to 128. However, samples are “reused,” which is acceptable when random sampling is used with a sufficient number of repetitions (*N* = 1000 in our application). The scheme without replacement avoids sample reuse, but we can have only a limited number of repetitions (average ∼226–310; Table S2B) and, more seriously, a much smaller maximum MUS size (16, rather than 128 with replacement). Given these differences, it is impossible for both schemes to achieve exactly the same results. In fact, since random sampling is used, even for the same scale size, results can differ within the same sampling scheme. This explains the minor differences in model parameters between the unit scale (US) (Table 2) and those for MUS = 1 (Table S2).

Since there is no perfect solution to reconcile potential differences between the two schemes, we use the with-replacement scheme primarily to demonstrate our methods, as previously (Tables and Supplementary Tables numbered with A, such as Table 2A, Table S2A). Importantly, the issue is insignificant: results from the without-replacement scheme (B) are consistent in terms of patterns and trends with the previously exposed results.

## 4. Conclusions and Discussion

### 4.1 Conclusions

We have integrated three families of power laws — Taylor’s power law (TPL) for ecological heterogeneity, Smith’s power law (SPL) for environmental heterogeneity, and the diversity power laws (DAR, DMR, DVR, DHR) for diversity scaling — into a unified framework for analyzing the diversity–heterogeneity nexus in microbial ecosystems. By systematically varying two orthogonal factors — scale (unit scale vs. multi-unit scale) and accrual (without vs. with sample accumulation) — we have revealed several key insights.

**First, ecological heterogeneity at community scale shows mixed habitat-dependence** depending on the sampling scheme. For classic TPL (community scale, without accrual), significant differences between gut and vaginal were observed under with-replacement (Table S7A; *p* = 0.000), but these became non-significant under without-replacement (Table S7B; p=0.184). For accrual TPL (metacommunity scale, with accrual), exponents were significantly different under both schemes (*p* = 0.000).

For SPL, which scales **environmental heterogeneity** and also operates on metacommunity scale, the Q-V variant showed significant differences in all parameters under both schemes (*p* ≤ 0.001), while the Index-V variant showed mixed results: with replacement, ln(a) and HCT were non-significant (*p* = 0.497 and 0.324), but ln(a) became significant without replacement (*p* = 0.017 and 0.000).

**Second, scale affects heterogeneity scaling in distinct ways**. At MUS scale, accrual TPL is scale-invariant for AGP with replacement, while classic TPL remains scale-dependent. Other scale invariance patterns show mixed results. Although scale invariance is an attractive feature for predicting critical transitions such as microbiome dysbiosis, we caution that practical microbiome data may not support this theory, as only approximately 17% of the power law models we tested demonstrated rigorous scale-invariance.

**Third, diversity scaling differs qualitatively between habitats**, with virtually all diversity scaling parameters showing significant differences between AGP and vaginal microbiomes. The gut microbiome exhibits positive *z* at *q* = 0 (species richness) but negative *z* at *q* = 2, 3 (dominant species), indicating that as sampling effort increases, rare species accumulate while dominant species diversity declines. The vaginal microbiome maintains positive (though weak) *z* across all orders, reflecting its low richness and high dominance by *Lactobacillus*.

#### Fourth, the diversity–heterogeneity nexus follows predictable power laws in majority models

At *q* = 0, diversity and heterogeneity are tightly coupled; at higher orders, the relationship may reverse. TPL provides a tool for relating ecological heterogeneity and diversity, while SPL provides a tool for relating environmental heterogeneity and diversity, supporting the reciprocity principle (Ma & Ellison, 2026). Among the DPL models, DHR (*H* = *V*/*M*) is most aligned with the diversity–heterogeneity nexus and is recommended for future studies. The three integrated power law families constitute a triple power law approach for scaling diversity and heterogeneity, which we further discuss before concluding this article.

**Finally, the triple power law framework — integrating TPL, SPL, and DPL** — extends the original dual framework of TPL and SPL (agents and template, ecological and environmental heterogeneity; Ma *et al.,* 2026) beyond microbial ecology. The same power law relationships may govern any complex system where structured variation arises from interacting components, from ecosystems to economies to artificial intelligence.

### 4.2 The Diversity–Heterogeneity Nexus

The fourth conclusion summarized above states that the diversity–heterogeneity nexus follows predictable power laws, with TPL and SPL providing complementary tools for relating heterogeneity and diversity, and DHR being the most aligned diversity-scaling model. Tables 4–6 present linear regression models relating corresponding parameters (e.g., *b* of TPL vs. *b* of SPL; *z* of DAR vs. *z* of DMR; *b* of TPL vs. *z* of DPL). The first parameter in each table is the slope (β), and the second is the intercept (α) of the linear function.

**Table 6.** Parameters of linear regression models relating the *b*-values of classic TPL, accrual TPL, SPL (V-i), and SPL (V-Q) to the z-values of DAR (D-Q), DMR, DVR, and DHR, with microbiome sites (AGP or vaginal) and sampling schemes (with/without replacement) all modeled separately (four regimes in total). This is an excerpted version (see Table S8 for the full version including *q* = 1 − 4).

| Orders | Correlations | AGP | | | | Vaginal Microbiome | | | | $N$ |
| --- | --- | --- | --- | --- | --- | --- | --- | --- | --- | --- |
| | | $\beta$ | $\alpha$ | $R$ | $p$ | $\beta$ | $\alpha$ | $R$ | $p$ | |
| **With Replacement** |  |  |  |  |  |  |  |  |  |  |
| $q=0$ | Classic TPL vs. DAR(Q) | 0.365 | -0.643 | 0.525 | 0.000 | -0.079 | 0.309 | 0.041 | 0.647 | 128 |
|  | Classic TPL vs. DMR | 0.361 | -0.635 | 0.510 | 0.000 | -0.068 | 0.284 | 0.035 | 0.691 | 128 |
|  | Classic TPL vs. DVR | 0.181 | -0.319 | 0.516 | 0.000 | -0.048 | 0.174 | 0.049 | 0.585 | 128 |
|  | Classic TPL vs. DHR | 0.365 | -0.642 | 0.522 | 0.000 | -0.123 | 0.409 | 0.061 | 0.497 | 128 |
|  | SPL(i) vs. DAR(Q) | -0.623 | 1.295 | 0.287 | 0.001 | -2.002 | 4.119 | 0.382 | 0.000 | 128 |
|  | SPL(i) vs. DMR | -0.740 | 1.529 | 0.335 | 0.000 | -2.073 | 4.261 | 0.399 | 0.000 | 128 |
|  | SPL(i) vs. DVR | -0.357 | 0.738 | 0.325 | 0.000 | -1.110 | 2.278 | 0.416 | 0.000 | 128 |
|  | SPL(i) vs. DHR | -0.688 | 1.425 | 0.316 | 0.000 | -2.378 | 4.870 | 0.434 | 0.000 | 128 |
|  | SPL(Q) vs. DAR(Q) | -0.623 | 1.295 | 0.287 | 0.001 | -2.002 | 4.119 | 0.382 | 0.000 | 128 |
|  | SPL(Q) vs. DMR | -0.740 | 1.529 | 0.335 | 0.000 | -2.073 | 4.261 | 0.399 | 0.000 | 128 |
|  | SPL(Q) vs. DVR | -0.357 | 0.738 | 0.325 | 0.000 | -1.110 | 2.278 | 0.416 | 0.000 | 128 |
|  | SPL(Q) vs. DHR | -0.688 | 1.425 | 0.316 | 0.000 | -2.378 | 4.870 | 0.434 | 0.000 | 128 |
|  | Accrual TPL vs. DAR(Q) | 0.038 | -0.024 | 0.007 | 0.935 | -4.478 | 9.048 | 0.678 | 0.000 | 128 |
|  | Accrual TPL vs. DMR | 0.216 | -0.380 | 0.041 | 0.644 | -4.469 | 9.030 | 0.684 | 0.000 | 128 |
|  | Accrual TPL vs. DVR | 0.073 | -0.120 | 0.028 | 0.754 | -2.355 | 4.756 | 0.701 | 0.000 | 128 |
|  | Accrual TPL vs. DHR | 0.078 | -0.104 | 0.015 | 0.866 | -4.953 | 9.995 | 0.718 | 0.000 | 128 |
|  | Accrual TPL vs. DHR | -0.991 | 1.982 | 0.999 | 0.000 | -1.026 | 2.051 | 0.996 | 0.000 | 128 |
| $q=1$ | ... | | | | | | | | | |
| $q=2$ | ... | | | | | | | | | |
| $q=3$ | Classic TPL vs. DAR(Q) | 0.024 | -0.046 | 0.158 | 0.076 | -0.071 | 0.165 | 0.233 | 0.008 | 128 |
|  | Classic TPL vs. DMR | 0.026 | -0.051 | 0.157 | 0.077 | -0.073 | 0.170 | 0.238 | 0.007 | 128 |
|  | Classic TPL vs. DVR | 0.013 | -0.025 | 0.155 | 0.081 | -0.036 | 0.085 | 0.232 | 0.008 | 128 |
|  | Classic TPL vs. DHR | 0.025 | -0.048 | 0.152 | 0.087 | -0.071 | 0.167 | 0.224 | 0.011 | 128 |
|  | SPL(i) vs. DAR(Q) | 0.179 | -0.358 | 0.375 | 0.000 | -0.664 | 1.330 | 0.812 | 0.000 | 128 |
|  | SPL(i) vs. DMR | 0.230 | -0.459 | 0.438 | 0.000 | -0.670 | 1.342 | 0.811 | 0.000 | 128 |
|  | SPL(i) vs. DVR | 0.112 | -0.224 | 0.431 | 0.000 | -0.347 | 0.695 | 0.823 | 0.000 | 128 |
|  | SPL(i) vs. DHR | 0.219 | -0.437 | 0.424 | 0.000 | -0.718 | 1.436 | 0.834 | 0.000 | 128 |
|  | SPL(Q) vs. DAR(Q) | 0.179 | -0.358 | 0.375 | 0.000 | -0.664 | 1.330 | 0.812 | 0.000 | 128 |
|  | SPL(Q) vs. DMR | 0.230 | -0.459 | 0.438 | 0.000 | -0.670 | 1.342 | 0.811 | 0.000 | 128 |
|  | SPL(Q) vs. DVR | 0.112 | -0.224 | 0.431 | 0.000 | -0.347 | 0.695 | 0.823 | 0.000 | 128 |
|  | SPL(Q) vs. DHR | 0.219 | -0.437 | 0.424 | 0.000 | -0.718 | 1.436 | 0.834 | 0.000 | 128 |
|  | Accrual TPL vs. DAR(Q) | -1.085 | 2.169 | 0.957 | 0.000 | -0.995 | 1.987 | 0.966 | 0.000 | 128 |
|  | Accrual TPL vs. DMR | -1.170 | 2.338 | 0.941 | 0.000 | -1.007 | 2.012 | 0.968 | 0.000 | 128 |
|  | Accrual TPL vs. DVR | -0.580 | 1.159 | 0.940 | 0.000 | -0.511 | 1.021 | 0.962 | 0.000 | 128 |
|  | Accrual TPL vs. DHR | -1.149 | 2.295 | 0.939 | 0.000 | -1.033 | 2.063 | 0.953 | 0.000 | 128 |
| **Without Replacement** |  |  |  |  |  |  |  |  |  |  |
| $q=0$ | Classic TPL vs. DAR(Q) | -0.051 | 0.391 | 0.634 | 0.008 | 0.019 | 0.768 | 0.369 | 0.159 | 16 |
|  | Classic TPL vs. DMR | -0.050 | 0.387 | 0.638 | 0.008 | 0.024 | 0.815 | 0.334 | 0.206 | 16 |
|  | Classic TPL vs. DVR | -0.030 | 0.205 | 0.648 | 0.007 | 0.023 | 0.408 | 0.578 | 0.019 | 16 |
|  | Classic TPL vs. DHR | -0.068 | 0.428 | 0.659 | 0.006 | 0.072 | 0.808 | 0.521 | 0.039 | 16 |
|  | SPL(i) vs. DAR(Q) | -0.567 | 1.411 | 0.956 | 0.000 | -0.159 | 1.085 | 0.231 | 0.390 | 16 |
|  | SPL(i) vs. DMR | -0.554 | 1.384 | 0.967 | 0.000 | -0.671 | 2.043 | 0.697 | 0.003 | 16 |
|  | SPL(i) vs. DVR | -0.330 | 0.799 | 0.971 | 0.000 | -0.329 | 1.033 | 0.617 | 0.011 | 16 |
|  | SPL(i) vs. DHR | -0.732 | 1.743 | 0.973 | 0.000 | -0.604 | 2.015 | 0.325 | 0.220 | 16 |
|  | SPL(Q) vs. DAR(Q) | -0.567 | 1.411 | 0.956 | 0.000 | -0.159 | 1.085 | 0.231 | 0.390 | 16 |
|  | SPL(Q) vs. DMR | -0.554 | 1.384 | 0.967 | 0.000 | -0.671 | 2.043 | 0.697 | 0.003 | 16 |
|  | SPL(Q) vs. DVR | -0.330 | 0.799 | 0.971 | 0.000 | -0.329 | 1.033 | 0.617 | 0.011 | 16 |
|  | SPL(Q) vs. DHR | -0.732 | 1.743 | 0.973 | 0.000 | -0.604 | 2.015 | 0.325 | 0.220 | 16 |
|  | Accrual TPL vs. DAR(Q) | -0.484 | 1.242 | 0.990 | 0.000 | -0.229 | 1.239 | 0.648 | 0.007 | 16 |
|  | Accrual TPL vs. DMR | -0.466 | 1.206 | 0.988 | 0.000 | 0.190 | 0.505 | 0.384 | 0.142 | 16 |
|  | Accrual TPL vs. DVR | -0.278 | 0.693 | 0.992 | 0.000 | -0.145 | 0.730 | 0.530 | 0.035 | 16 |
|  | Accrual TPL vs. DHR | -0.616 | 1.508 | 0.993 | 0.000 | -0.832 | 2.530 | 0.872 | 0.000 | 16 |
| $q=1$ | ... .. | | | | | | | | | |
| $q=2$ | ... .. | | | | | | | | | |
| $q=3$ | Classic TPL vs. DAR(Q) | -0.094 | 0.209 | 0.672 | 0.004 | 0.038 | -0.018 | 0.271 | 0.310 | 16 |
|  | Classic TPL vs. DMR | -0.095 | 0.210 | 0.665 | 0.005 | 0.043 | -0.022 | 0.273 | 0.306 | 16 |
|  | Classic TPL vs. DVR | -0.045 | 0.099 | 0.676 | 0.004 | 0.023 | -0.015 | 0.283 | 0.288 | 16 |
|  | Classic TPL vs. DHR | -0.083 | 0.183 | 0.688 | 0.003 | 0.050 | -0.039 | 0.294 | 0.269 | 16 |
|  | SPL(i) vs. DAR(Q) | -0.985 | 1.972 | 0.963 | 0.000 | 0.079 | -0.080 | 0.042 | 0.877 | 16 |
|  | SPL(i) vs. DMR | -1.005 | 2.010 | 0.960 | 0.000 | 0.076 | -0.068 | 0.036 | 0.895 | 16 |
|  | SPL(i) vs. DVR | -0.467 | 0.934 | 0.962 | 0.000 | 0.052 | -0.059 | 0.047 | 0.864 | 16 |
|  | SPL(i) vs. DHR | -0.850 | 1.701 | 0.962 | 0.000 | 0.139 | -0.180 | 0.060 | 0.824 | 16 |
|  | SPL(Q) vs. DAR(Q) | -0.985 | 1.972 | 0.963 | 0.000 | 0.079 | -0.080 | 0.042 | 0.877 | 16 |
|  | SPL(Q) vs. DMR | -1.005 | 2.010 | 0.960 | 0.000 | 0.076 | -0.068 | 0.036 | 0.895 | 16 |
|  | SPL(Q) vs. DVR | -0.467 | 0.934 | 0.962 | 0.000 | 0.052 | -0.059 | 0.047 | 0.864 | 16 |
|  | SPL(Q) vs. DHR | -0.850 | 1.701 | 0.962 | 0.000 | 0.139 | -0.180 | 0.060 | 0.824 | 16 |
|  | Accrual TPL vs. DAR(Q) | -0.841 | 1.679 | 0.998 | 0.000 | -0.963 | 1.883 | 0.992 | 0.000 | 16 |
|  | Accrual TPL vs. DMR | -0.860 | 1.717 | 0.998 | 0.000 | -1.073 | 2.097 | 0.993 | 0.000 | 16 |
|  | Accrual TPL vs. DVR | -0.399 | 0.796 | 0.996 | 0.000 | -0.563 | 1.099 | 0.991 | 0.000 | 16 |
|  | Accrual TPL vs. DHR | -0.723 | 1.443 | 0.993 | 0.000 | -1.163 | 2.266 | 0.987 | 0.000 | 16 |

#### 4.2.1 TPL and SPL as Complementary Tools

Table 2 shows that classic TPL and SPL capture distinct aspects of heterogeneity. Under with-replacement sampling, classic TPL showed weak or non-significant linear relationships with SPL variants ( R ≈ 0.002, *p* = 0.980), confirming that cross-sectional ecological heterogeneity (classic TPL) and environmental heterogeneity (SPL) are not interchangeable. However, SPL showed moderate to strong linear relationships with accrual TPL (R ≈ 0.25 − 0.79, *p* = 0.005), supporting the reciprocity principle (Ma & Ellison, 2026): environmental heterogeneity and ecological heterogeneity are related through cumulative (accrual) processes. The slopes and intercepts of these linear regressions quantify how changes in one heterogeneity parameter predict changes in the other.

#### 4.2.2 DPL Redundancy and the Special Role of DHR

Table 3 demonstrates that the four DPL models are highly redundant. With-replacement sampling yielded near-perfect linear relationships among all DPL models across all diversity orders, confirming that DAR, DMR, DVR, and DHR carry essentially identical information about diversity scaling. The slopes are close to 1 and intercepts near 0, indicating that the scaling exponents of these models are nearly proportional. Without-replacement sampling produced similar patterns, though with slightly lower correlations for two models of vaginal at *q*=0. This redundancy suggests that researchers may not need to fit all four models; DAR, being the simplest, may suffice for most applications. However, DHR holds a special place: it uses *H* = *V*/*M*, a direct measure of heterogeneity, making it the most natural bridge between diversity scaling and heterogeneity scaling. For studies focused on the diversity–heterogeneity nexus, DHR is the preferred model.

#### 4.2.3 Heterogeneity Scaling Predicts Diversity Scaling in Majority Models

Table 4 confirms that heterogeneity scaling predicts diversity scaling in majority cases (∼60% in Table 4, and ∼70% in the full results in Table S8), with the nature of this prediction depending on the heterogeneity model and diversity order. Classic TPL showed weak to moderate linear relationships with DPL at q=0 in AGP, but weaker relationships in vaginal, suggesting that cross-sectional ecological heterogeneity primarily relates to species richness scaling. SPL variants showed moderate linear relationships with DPL in vaginal, supporting the reciprocity principle: environmental heterogeneity is particularly important for diversity scaling in low-diversity, dominance-driven habitats. Most strikingly, accrual TPL showed strong negative linear relationships with DPL at high *q* in both habitats (R ≈ 0.95, *p* = 0.000), with slopes indicating that as heterogeneity accumulation increases, higher-order diversity scaling decreases—a stabilization effect.

### 4.3 Integration with Existing TPL and SPL Literature

The literature on Taylor’s power law (TPL) is vast, spanning disciplines from ecology and agriculture to physics, economics, finance, and even the humanities (Eisler et al., 2008; Meng 2015, Ma & Taylor, 2025). Meng (2015) examined five decades of TPL studies, clarifying that Smith (1938) investigated the relationship between crop yield variance and plot size—not the variance–mean relationship that defines TPL. Eisler et al. (2008), in their influential review, mistakenly conflated Smith’s law with TPL, but their contribution remains significant for emphasizing the temporal dynamics of TPL and its connections to fluctuation scaling in physical systems—a perspective extended by De La Pena et al. (2022) to dependent samples.

Ma & Taylor (2025) extended the review to six decades, identifying three distinct periods and eight major themes spanning ecological mechanisms, statistical distributions, mathematical foundations, population stability, tipping-point signals, and applications in complex networks and microbiomes. Recent work on microbiome variability has shown that both spatiotemporal noise and macroecological dynamics can be quantified through scaling relationships (Ji et al., 2019; Ji et al., 2020; Ma, 2015, 2025), underscoring the relevance of power-law approaches for understanding microbial community structure.

In contrast, the literature on Smith’s power law (SPL) is comparatively sparse. Since Smith’s original 1938 paper, SPL has remained largely confined to agricultural and soil science. Recent efforts have begun to address this gap. Ma et al. (2026) proposed a dual framework integrating TPL and SPL, arguing that together they offer a more complete picture of heterogeneity scaling across ecological and environmental domains. The present article extends this dual framework to a triple power law framework by incorporating diversity power laws (DPL), thereby linking heterogeneity scaling with diversity scaling.

The evidence from Tables 4–6 supports this integration. TPL and SPL provide complementary tools for relating heterogeneity and diversity, with accrual TPL showing the strongest predictive relationship at higher diversity orders. Among the DPL models, DHR—using the variance-to-mean ratio as a direct heterogeneity metric—is most aligned with the diversity–heterogeneity nexus. Linear regression models confirm that the scaling parameters of these power laws are systematically related, providing a quantitative framework for the triple power law approach.

Nevertheless, predictive power appears effective only in a majority (∼60–70%) of models. This percentage, together with the ∼17% scale-invariance rate, reveals a sobering reality: pursuing neat and ideal power-law predictions can be challenging, even beyond the cautions raised by Stumpf & Porter (2012). The relationships are complex, and further investigations are needed. Our rigorous scale-invariance testing revealed that fewer than 20% of the models examined were strictly scale-invariant, underscoring the complexity of scaling relationships in real-world microbial ecosystems. This finding resonates with the long-standing tension in the TPL literature—whether it is a universal law or a scale-dependent empirical pattern—and suggests that the answer may depend on the specific model, scale range, and sampling scheme.

Beyond TPL and SPL, several related developments merit mention. Döring et al. (2015) proposed using power law residuals (POLAR) as a stability metric for crop yields. Kendal (2004) showed that Tweedie distributions—scale-invariant exponential dispersion models—can generate TPL, and Cohen & Xu (2015) demonstrated that TPL emerges from any skewed distribution with four finite moments. Cohen & Schuster (2012) proposed Variance-Mass Allometry (VMA), linking population variance to mean body mass. These developments underscore the broad applicability and diverse mathematical underpinnings of power laws in ecology and beyond.

The triple power law method is a simplified model of the diversity–heterogeneity nexus, while AGP and VM represent real-world data; a perfect fit is unlikely. An equally significant contribution of this article is providing a tool for detecting deviations between theoretical expectations and real-world data, extending beyond the predictive models themselves. Nevertheless, the empirical and theoretical development of SPL still lags far behind that of TPL.

### 4.4 Towards a Unified Principle across Disciplines

The TPL–SPL–DPL framework reveals a general principle: complex systems, whether ecological or computational, exhibit dual scaling laws for agents and their templates, with diversity as the emergent outcome. This principle finds precise parallels in modern computing and AI. Moore’s law (hardware scaling) and Koomey’s law (energy efficiency) capture the agent–template duality in physical systems; the OpenAI–DeepMind scaling debate—model size vs. data—captures the same tension in software. In both cases, optimizing the agent alone is insufficient; the template must scale with it, and the final performance (benchmark scores, energy efficiency, or biodiversity) and/or applications are the emergent outcome of their interaction (Table 5).

**Table 7.** Agent (ecological heterogeneity) and template (environmental heterogeneity) interactions generate emergent property scaling across ecology, hardware, and AI.

| Domain | Agent (TPL) | Template (SPL) | Outcome (DPL) |
| --- | --- | --- | --- |
| Ecosystems | Ecological heterogeneity of organisms by TPL (Taylor 1961) | Environmental (habitat) heterogeneity by SPL (Smith 1938) | Biodiversity by DPL (Watson 1838, Ma 2018), Ma et al. (2026) |
| Computation | Moore's (1965) law (compute) | Koomey's (2011) law (energy) | Benchmark scores, Energy Efficiency, Applications such as Web, Mobile APP, Games, ... .. |
| AI-LLMs | OpenAI scaling (model size) (Kaplan et al. 2020) | DeepMind scaling (data) Hoffmann et al. (2022) | Agents, Robots, ... |

In computer science, people recognized the importance of chips and energy from the very beginning—because they had to make it happen. Without hardware and energy, nothing runs. In ecology, nature made it happen. The environment is already there, and we often ignore its heterogeneity, jumping directly to measure diversity. We count species richness, but often ignore their interactions—which is the essence of heterogeneity. Diversity is a count; heterogeneity is a relationship. Counting species without understanding how they interact, compete, and cooperate misses the very structure that generates diversity in the first place. This unavoidable shortcut creates gaps and mishaps: we measure diversity without quantifying the environmental heterogeneity and the biotic interactions that sustain it. The template—whether silicon or soil—is the invisible half.

In computer science, the agent–template duality is foundational. Moore’s law (compute) and Koomey’s law (energy) represent the hardware-level scaling of agents and their templates. OpenAI’s scaling law (model size) (Kaplan et al. 2020) and DeepMind’s refinement (data) Hoffmann et al. (2022) represent the software-level counterpart. The applications that ultimately matter—web, mobile apps, social media, games—are the emergent outcomes of these scaling laws (chip density and energy efficiency). They are the “diversity” of the digital world: the tangible products that users interact with and value.

In our framework, DPL plays the same role. Biodiversity is not merely a byproduct of ecological and environmental heterogeneity; it is the outcome that human society cares most about—the living fabric of our planet. Just as applications are the realized value of computational scaling, biodiversity is the realized value of ecological scaling. The triple power law framework thus aligns ecology with computer science: both are stories of agents, templates, and the emergent diversity they generate together.

It is easy to forget the template. Moore’s law—the doubling of transistors on a chip every two years—is celebrated as the engine of computing progress (Moore 1965). Koomey’s (1965) law—the doubling of energy efficiency of computing every 1.5 years—is rarely mentioned, even though it is what makes modern computing physically sustainable (otherwise, your phone would become a portable heater, or worse a ticking time bomb). In ecology, Smith’s law remains largely unknown outside crop science—despite its foundational importance. The template is the invisible half. Just as ecological diversity depends on both organisms and their habitats, AI performance depends on both models and data. Rediscovering the template—whether SPL or Koomey’s law—offers a unified lens to predict heterogeneity and diversity across space, time, and disciplines, from ecosystems to economies to AI governance.

